# Evolution of a costly immunity to cestode parasites is a pyrrhic victory

**DOI:** 10.1101/2021.08.04.455160

**Authors:** Jesse N. Weber, Natalie C. Steinel, Foen Peng, Kum Chuan Shim, Brian K. Lohman, Lauren Fuess, Stephen de Lisle, Daniel I. Bolnick

## Abstract

Parasites impose fitness costs on their hosts. Biologists therefore tend to assume that natural selection favors infection-resistant hosts. Yet, when the immune response itself is costly, theory suggests selection may instead favor loss of resistance. Immune costs are rarely documented in nature, and there are few examples of adaptive loss of resistance. Here, we show that when marine threespine stickleback colonized freshwater lakes they gained resistance to the freshwater-associated tapeworm, *Schistocephalus solidus*. Extensive peritoneal fibrosis and inflammation contribute to suppression of cestode growth and viability, but also impose a substantial cost of reduced fecundity. Combining genetic mapping and population genomics, we find that the immune differences between tolerant and resistant populations arise from opposing selection in both populations acting, respectively, to reduce and increase resistance consistent with divergent optimization.

**One Sentence Summary:** Recently-evolved freshwater populations of stickleback frequently evolve increased resistance to tapeworms, involving extensive fibrosis that suppresses parasite growth; because this fibrosis greatly reduces fish fecundity, in some freshwater populations selection has favored an infection-tolerant strategy with fibrosis suppression.

## Main Text

Parasites are a major source of natural selection on their hosts, driving rapid evolution in host behavior, physiology, and especially immunity, to avoid or eliminate infections (Fumagalli et al. 2011; Ebert and Fields 2020; Little 2002). Parasites, in turn, evolve counter-measures that evade or suppress host immunity to establish long-term infections or to facilitate their transmission (Schmid-Hempel 2008; Maizels et al. 2004). A fundamental question in host-parasite biology is why one party does not ‘win’ this coevolutionary race (Nuismer 2017)? Why don’t hosts evolve fully effective immunity that drives the parasite to extinction? A potential answer to this question is that the costs of immunity might be too great to justify the evolution of perfectly resistant hosts (Boots and Bowers 2004; Tschirren and Richner 2006). Instead, hosts might evolve an intermediate level of immunity that optimizes the balance between marginal benefits versus costs, but which allows persistence of the parasite population (Urban, Bürger, and Bolnick 2013; Viney, Riley, and Buchanan 2005). Costs of immunity are well supported in laboratory studies of model organisms such as mice, and in livestock (Lochmiller and Deerenberg 2000; Hasselquist and Nilsson 2012; van der Most et al. 2011; Rauw 2012; Nystrand and Dowling 2020), but remain largely undocumented in wild organisms. Here, we present evidence of both benefits and costs of recently-evolved resistance by wild populations of a small fish (threespine stickleback) against a cestode parasite. Consistent with an optimization process, we observe positive selection acting on alleles that both increase and suppress sticklebacks’ immune response to infection.

### Population differences in infection rates

Like most animals, threespine stickleback (*Gasterosteus aculeatus*) carry a rich community of macroparasites. The composition and diversity of this community varies substantially among habitats (e.g., estuary, stream, or lake), and among populations of a given habitat (MacColl 2009; Bolnick et al. 2020). For example, the cestode *Schistocephalus solidus* predominantly infects lake populations of stickleback, but among lakes infection prevalence can range from 0% to over 75% (Fig. 1A; (Weber, Kalbe, et al. 2017)). Some of this variance in prevalence is driven by abiotic variables such as salinity because *S*.*solidus* eggs do not tolerate brackish water (Heins, Baker, and Green 2011). Accordingly, marine stickleback are rarely exposed to *S*.*solidus* (Fig. 1A), and have correspondingly weak immunity (∼90% infection rate in the lab), whereas more frequently-exposed lake populations evolved resistance (∼20% infection; (Weber, Kalbe, et al. 2017). However infection rates also vary among lakes. Some of this variance is ecological (e.g., lake elevation, Fig. S1; prey community Fig. S2), but even ecologically similar lakes may differ in infection rates. Stickleback from Roberts and Gosling Lakes (R and G hereafter) on Vancouver Island persistently differ in *S*.*solidus* prevalence (Fig. 1A; Weber,Steinel, et al. 2017), despite the lakes’ proximity (17 km apart) and similar size and elevation. Cyclopoid copepods, the first intermediate hosts for *S*.*solidus*, constitute a similar fraction (∼5%) of adult stickleback’s breeding-season diet in both lakes (Weber, Steinel, et al. 2017), and terminal hosts (loons and mergansers) breed in both lakes annually. We hypothesized that the persistently higher infection prevalence in G than R (Fig. 1A) reflects evolved differences in either resistance by R stickleback, or tolerance of G fish.

**Fig. 1.**
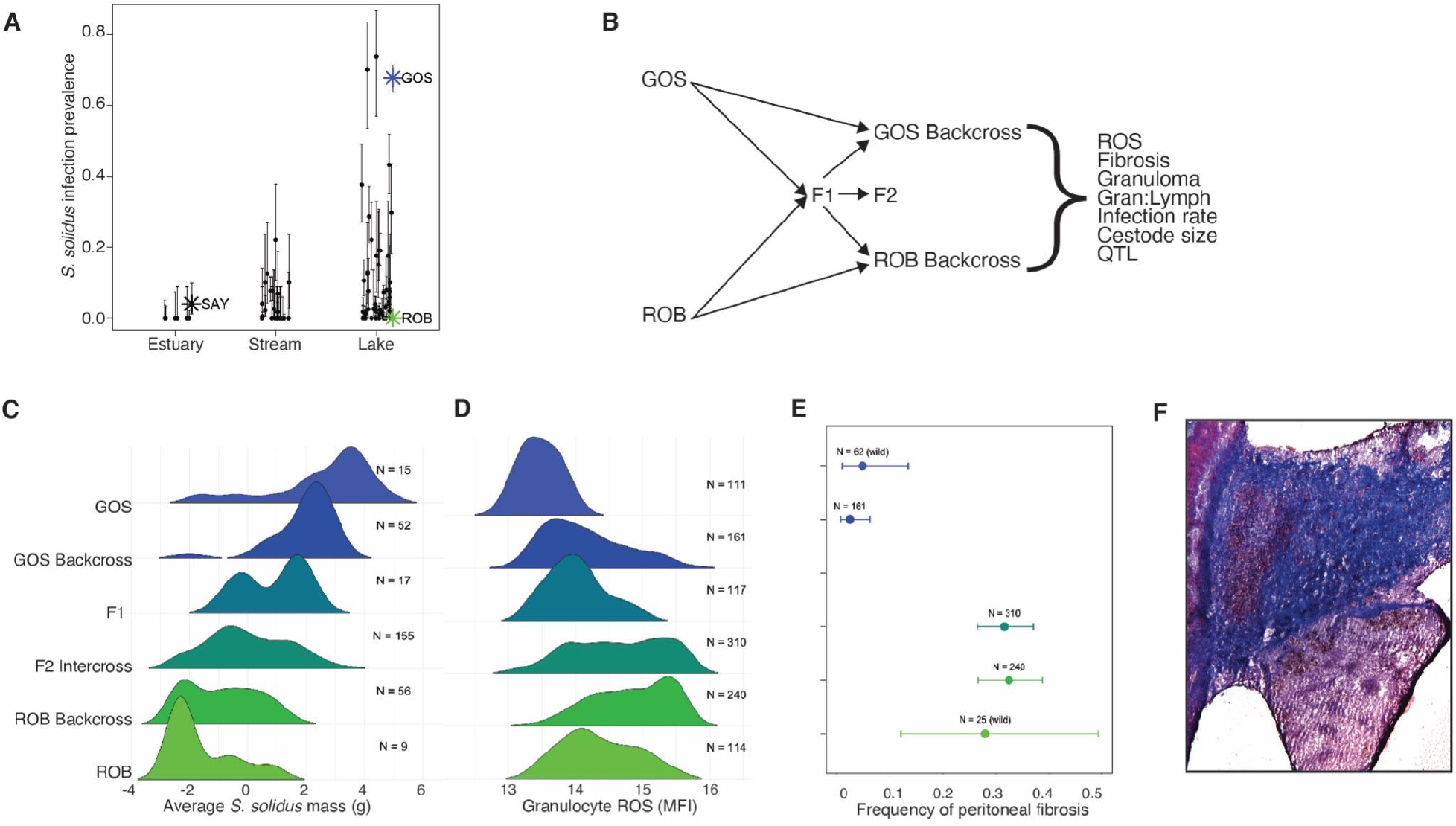
Population differences in infection success and immune phenotypes. **A)** The prevalence of *S*.*solidus* cestodes varies greatly between stickleback in different aquatic habitats on Vancouver Island, British Columbia (binomial GLM deviations = 148.8 df = 2 P < 0.0001), as well as among populations within a given habitat type. For instance prevalence varies significantly between lakes (c^2^ = 4629, df = 49, P < 0.0001; estimate and 95% CI plotted). Three focal populations are labeled: high-infection Gosling Lake (GOS), infection-free Roberts Lake (ROB) and a low-infection susceptible marine population from Sayward Estuary (SAY) representing the ancestral state. **B)** A diagram of the experimental design, infecting lab-reared F2 hybrids with S.solidus cestodes then assaying immune traits and infection outcomes for QTL mapping. **C**) Cestode mass is greatest in lab-infected Gosling Lake stickleback, least in Roberts Lake, and intermediate in F1, F2, and reciprocal backcross hybrids. **D)** Reactive oxygen production by granulocytes (mean fluorescent intensity, MFI, of gated granulocytes) differs between lab-raised GOS and ROB stickleback, and their F1 and F2 hybrids are intermediate (sample sizes listed for each cross time; MFI depends on cross type (F = 43.14, P < 0.0001), and log mass (F = 27.06, P < 0.0001) but not fish sex (F = 0.815, P = 0.4428) nor cestode presence (F = 2.06 P = 0.1518). A weak cross*Infection interaction (F = 2.47 P = 0.0308) exists because RBC backcross fish decrease ROS significantly when infected, whereas all other crosses exhibit no response. **E**) Peritoneal fibrosis is more common wild-caught ROB than GOS fish, and more common in lab-reared cestode-exposed RBC and F2 hybrids than GBC hybrids (Cross c^2^ = 155 P < 0.0001, sex c^2^ = 2.53 P = 0.281, log mass c^2^ = 23.83 P < 0.0001, cestode infection c^2^ = 98.48 P < 0.0001, cross*infection c^2^ =18.45 P = 0.002). **F**) Trichrome stain of a section through stickleback intestine and spleen showing fibrosis (purple-stained collagen) connecting the intestinal wall (left side) to the spleen (lower right).

### Heritable divergence in immune traits

We bred lab-reared R×G second generation intercross (hereafter noted as “F2”) hybrids and reciprocal backcrosses (RBC and GBC), and experimentally exposed the three cross types to *S*.*solidus* to genetically map heritable differences in infection outcomes and immune traits (Fig. 1B). In F2 hybrids greater R ancestry conferred a lower infection rate (Z = -2.005, P = 0.045; Fig. S3), consistent with a non-significant trend previously documented in pure R and G lab infections (Weber, Kalbe, et al. 2017; Weber, Steinel, et al. 2017). However, when cestodes succeeded in initiating infection, they grew slower in fish with more R ancestry: slower in R and RBC fish than in F2s, and slower in F2s than GBCs, and fastest in pure G and susceptible marine fish (Fig. 1C, proportion R ancestry t=-11.6, P < 0.0001). This analysis accounted for covariates including a weak tendency for cestodes to be larger in males (sex t=-2.5, P=0.013), in larger fish (ln fish mass t=3.9, P=0.001), and no significant effect of coinfection intensity (t=-1.0, P=0.308). *S*.*solidus’* life cycle requires transmission to a piscivorous bird terminal host, so cestode infections tend to not be directly fatal. Instead, as cestodes grow large enough to reproduce, they begin manipulating host behavior and physiology to facilitate predation. The cestode growth suppression by all R and RBC and many F2 fish prevented cestodes from reaching the minimum reproductive size of 50mg (Tierney and Crompton 1992), so should mitigate the cestode’s effect on host predation susceptibility.

Consistent with the difference in infection success, lab-raised R and G fish exhibit substantial differences in immune traits. Infection induced elevated granulocyte abundance in the hematopoietic organ in both host genotypes (Fig. S4), and in R fish these granulocytes generated more reactive oxygen species (ROS) per cell (Fig. 1D). In addition to the more intense inflammatory response in R fish, they developed severe peritoneal fibrosis following cestode exposure (Mittal et al. 2014); Fig. 1E). The peritoneum, a single layer of mesothelium supported by a vascularized connective tissue that lines the abdominal cavity secretes a lubricating serous fluid that facilitates the free movement of the visceral organs. *S. solidus* migrate out of the stickleback gut and through the peritoneum 12-24 hours after ingestion (Hammerschmidt and Kurtz 2007). Damage within the peritoneal cavity (e.g., the coelom) causes inflammation, leading to fibroblast proliferation, collagen deposition, and the formation of adhesions between the viscera and/or between the viscera and the abdominal wall (Buckman 1976). In lab-reared fish the organs move freely, but following cestode exposure R genotypes exhibit extensive adhesion among organs (Supplementary Video 1). We can induce similar adhesions in lab-reared fish by intraperitoneal injection of aluminum phosphate (a common vaccine adjuvant), or cestode protein extracts (Hund et al. 2020). Trichrome stain of histological sections from alum-injected fish confirmed these adhesions contain excessive collagen deposition consistent with fibrosis (Fig. 1F, Fig. S5). In our cestode-exposed F2 hybrids, fibrosis is more common in fish with more R ancestry (Fig. 1E) when a cestode is observed (Fig. S6A; binomial GLM analysis: host genotype effect, χ^2^=155.0, df = 1, P < 1e^-5^; infection effect, χ^2^=96.5, df = 1, P < 1e^-5^; genotype*infection interaction χ^2^=18.45, df = 1, P = 0.00243). Fibrosis is rare in G fish (in the wild and in the lab), and in lab GBC fish (Fig. 1E) regardless of infection. Covariates in the GLM indicate that fibrosis was more likely in larger individuals (χ^2^=23.8, df = 1, P < 1e^-5^), with no sex effect (χ^2^=2.53, df = 1, P = 0.28). Fibrosis is absent in marine stickleback (in the wild, and after laboratory infections), so its presence in some (but not all) post-Pleistocene lake populations represents an evolutionarily derived trait which has evolved numerous times in parallel (Fig. S7).

### Benefits of recently evolved immune traits

Fibrosis contributes to cestode resistance in two ways. First, cestodes in fibrotic F2 hosts were 87.7% smaller than the cestodes in non-fibrotic F2s (Fig 2A, mean mass 0.316g and 2.558g respectively, t=6.21, df = 156, P < 0.0001). This cestode growth suppression was also observed in RBC and GBC families, but was non-significant because nearly all cestodes were very small in RBC, and few GBC hybrids had fibrosis. Overall, cestode mass depended on both fibrosis (F_1,255_=7.32, P=0.0072) and cross (F_2,255_ = 56.2, P < 0.0001), but no fibrosis*cross interaction (F_2,255_= 1.25, P = 0.2873). Higher ROS was also associated with cestode growth suppression. The trend is only significant in F2 hybrid crosses (where variance in cestode size is greatest), yet the effect size is comparable in all three cross types (Fig 2B; main effect of ROS F_1,236_ = 5.25, P=0.0228 in a model also including fibrosis [F = 13.18, P< 0.001], cross [F = 37.9, P < 0.001]; P>0.05 for all interaction effects).

**Fig. 2.**
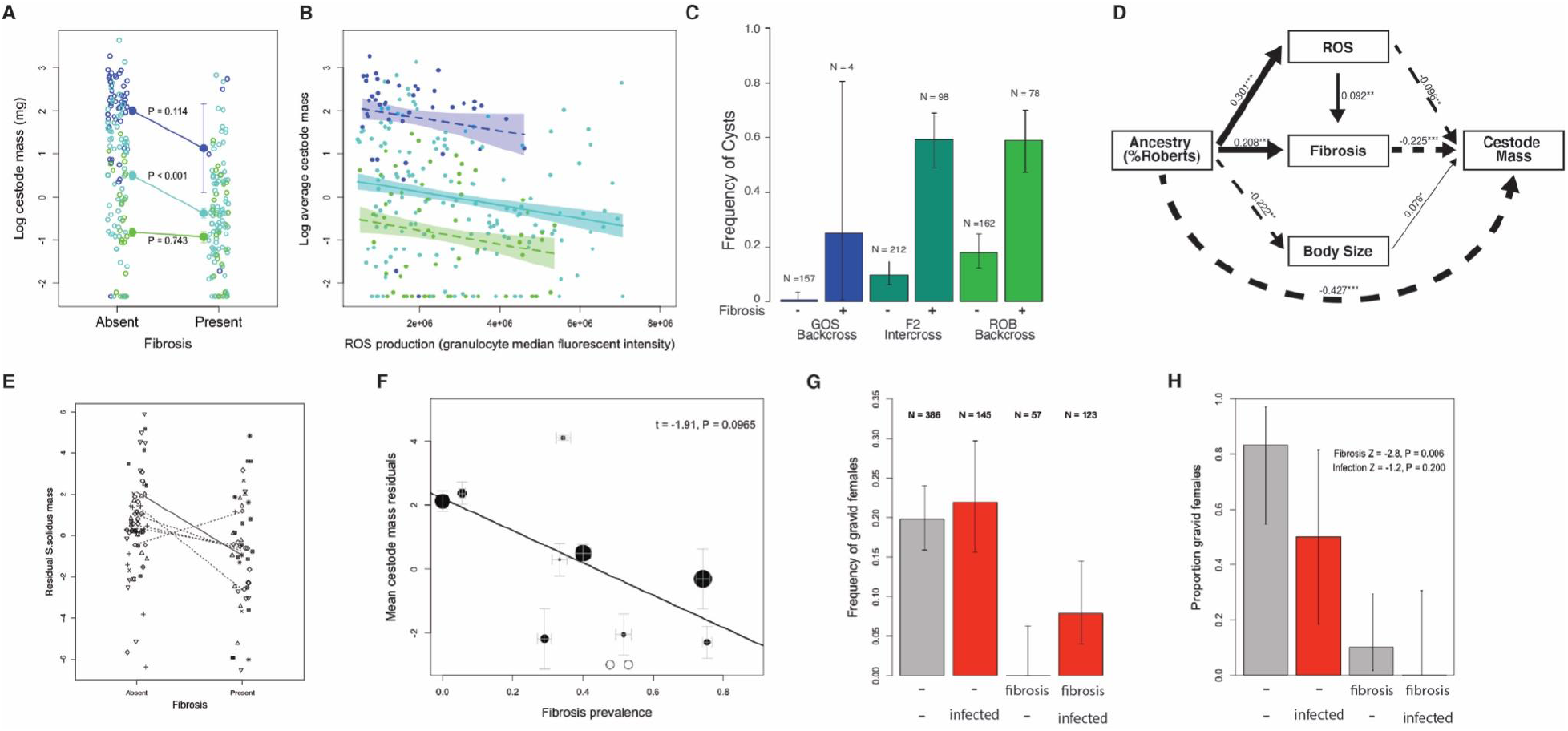
Benefits and costs of fibrosis and ROS in lab-infected stickleback. **A.** Cestode mass is lower in fish with greater R ancestry (proportion R ancestry t = -11.6 P < 0.0001; sex t = -2.505 P = 0.0127; log mass t = 3.86 P = 0.0014; number of coinfecting cestodes t = -1.02 P = 0.3082). Sample sizes listed to the right are for infected fish. Adding fibrosis as a term in this model does not appreciably change significance of the other terms, but confirms a significant negative effect of fibrosis on cestode mass (t = -2.909, P = 0.0039). **B.** Stickleback with greater ROS production have reduced cestode mass, within each of the three F2 hybrid cross categories, significantly in F2 intercrosses (solid trendline), with the same slope but non-significant in backcrosses (dashed trendline). **C.** Stickleback with fibrosis are more likely to have cysts encasing tapeworms (pictured in Fig. S7), particularly in F2 and RBC fish. **D.** Path analysis of the relationship between % ROB ancestry, fish mass, ROS production, fibrosis, and cestode mass. Dashed lines denote negative correlations, solid lines are positive. Thick lines are significant, thin lines are non-significant. **E.** Consistent with the laboratory infection result, on average wild-caught individuals with fibrosis have lower cestode mass than fibrosis-free individuals. Each trendline is a separate population (with different data point symbols), solid and dashed lines represent significant and non-significant within-lake trends, the main effect of fibrosis is significant in a model with random population effects. **F.** On Vancouver Island, lake populations with more prevalent fibrosis have a marginally non-significant tendency to have smaller cestodes (controlling for host mass). Fibrosis, but not cestode infection, were significantly associated with reduced reproductive development of lab-raised females. **G.** Costs of fibrosis are evident because lab-raised female stickleback are less likely to be reproductively mature (gravid) with fibrosis, compared to without, controlling for cestode presence. **H.** Similar costs to female fecundity are observed in wild-caught fish, here showing the result for Roselle Lake.

Second, stickleback with fibrosis are more likely to encase small cestodes in cysts, likely to be granulomas (Fig. S8), which often contain dead or visibly degraded parasites. RNAseq of ten cysts yielded overwhelmingly *S*.*solidus* transcripts (91.3% of 272 million reads mapped to the cestode genome, compared to 0.3% mapping to the stickleback genome), even when no identifiable cestode body remained. Having fibrosis significantly increased the likelihood of a cyst in RBC fish (odds ratio = 6.6, P < 0.0001), F2 fish (odds ratio = 13.2, P < 0.0001), and even in the few affected GBC fish (odds ratio = 52, P = 0.0359). Like fibrosis, cysts were significantly more common in fish with greater R ancestry (Fig. 2C; χ^2^=83.9, df = 5, P < 1e^-5^), or when there was a cestode present (χ^2^=31.8, df = 1, P < 1e^-5^). From these results we conclude that fibrosis is associated with growth suppression and the encysting and killing of small cestodes. Importantly, fibrosis is also persistent lesion that remains after even a transient immune challenge (e.g., >3 months after injection with aluminum phosphate or cestode protein extracts; (Hund et al.2020)), explaining why we observe cestode-exposed fish with residual fibrosis but no obvious parasites.

Path analysis suggests that the negative effect of R ancestry on cestode growth arises via multiple direct and indirect paths that account for 34.3% of cestode size variation (Fig. 2D): R ancestry is associated with both more fibrosis (r = 0.208, P = 0.001) and more ROS (r = 0.301, P < 0.001), but ROS and fibrosis are decoupled (r = 0.075, P = 0.242). R ancestry also confers smaller fish size (r = -0.222 P < 0.001), but this reduction in host size does not contribute to the cestode growth suppression (r = 0.076 P = 0.148). Smaller cestode mass is attributed to the fibrosis (r=-0.226, P < 0.001), but there remains a sizeable direct negative effect of R ancestry on cestode mass that is unexplained (r=-0.427, P<0.001; Fig. 2D).

Our lab results are confirmed by multiple independent samples of wild stickleback from lakes in British Columbia. In a sample of 13 lakes with significant variation in fibrosis prevalence (𝒳^2^=90.8, P<0.0001), cestode-infected individuals are more likely to have fibrosis (𝒳^2^ =4.4, P=0.034; Fig. S9). In contrast, another helminth infecting the peritoneum, *Eustronglyides sp*., had no association with fibrosis (𝒳^2^=0.98, P=0.197). We also observe fibrosis-related growth suppression in the wild: infected individuals with fibrosis had 56.6% lower cestode mass compared to non-fibrotic infected individuals (Fig. 2E; 𝒳^2^ = 20.9, df = 1, P = 0.0000047; host mass 𝒳^2^ = 0.075, df = 1, P = 0.7833, with lake random effect). Comparing across another sample of 16 lakes (Stuart et al 2017), cestode prevalence was positively correlated with fibrosis (r=0.580, P=0.0009). In that survey, cestode mass was marginally (non-significantly) smaller in populations with more fibrosis (Fig. 2F; t = -1.91, P = 0.0965, controlling for crowding infection intensity). These field observations match our lab finding that cestode infection tends to induce fibrosis, which then acts to reduce cestode mass for individual hosts, and among populations.

### Costs of recently-evolved immune traits

Both fibrosis and ROS are costly responses to infection. In the laboratory-exposed hybrid fish, fibrosis was associated with a 73.4% reduction in the frequency of gravid females, from 20.4% of females with mature ovaries, to 5.4% at the time of euthanasia (Fig. 2G). Overall, a binomial GLM indicates that females’ reproductive maturity improved with fish log mass (deviance = 30.1 P < 0.0001), and decreased with both fibrosis (d = 13.53, P = 0.0002, Fig. 2G) and ROS (d = 16.14 P < 0.0001, Fig. S10), but not cross type (d = 3.48 P = 0.175) or cestode infection (d=0.14, P = 0.703). The presence of a cyst had no detectable effect on reproductive state (P=0.137). From these results we can conclude that in its early stages, cestode infection undermines female fitness indirectly via fibrosis. Direct effects of infection might emerge later as cestodes continue to grow (Bagamian, Heins, and Baker 2004; T. Schultz, Topper, and C. Heins 2006). Field samples corroborate a central role of fibrosis in mediating the negative effect of *S*.*solius* infection on female fitness. Focusing on one of the few lakes with an intermediate frequency of both infection and fibrosis (Roselle Lake), female reproductive maturity was not significantly affected by infection (z = -1.283 P = 0.1996), but was reduced 90.5% by fibrosis (z = -2.7, P = 0.0057) from 70% to 6.6% (Fig. 2H). This trend in wild stickleback closely mirrored the effect sizes in our lab-raised study (Fig. 2G).

Comparable costs exist for male stickleback. If fibrosis reduces the ability or motivation to build a nest and defend it against rivals and egg predators, we expect fibrosis rates of successfully nesting males to be lower than non-nesting males caught nearby outside of the aggregation of nests. In each of two lakes (Roselle and Boot) we confirmed that nesting males had lower fibrosis than males who had failed to nest (odds ratios of 0.41 and 0.56 respectively, both P < 0.01), controlling for infection status and body size (De Lisle and Bolnick 2020). The mechanisms of these fitness costs of fibrosis are still under investigation, but may include increased metabolic demands, locomotor constraints from more rigid tissues, or constrained female ovary expansion due to adhesion to surrounding tissue.

### Genetic basis of variation in cestode resistance

The benefits and costs of fibrosis, described above, lead us to hypothesize that fibrosis may be subject to stabilizing selection with optima that differ depending on local conditions (e.g., exposure risk, severity of costs). Such optimization should be reflected in the genetic architecture of the relevant traits, with selection acting on a combination of pro-and anti-fibrotic genes (positive and negative effects) rather than systematic directional selection for ever-higher fibrosis. To test this expectation, we next sought to identify the genetic basis of this immune variation by triangulating a combination of QTL mapping, whole genome sequencing, and transcriptomics.

To locate chromosomal regions contributing to G-R phenotypic differences we experimentally exposed 647 F2, GBC and RBC fish to *S*.*solidus*, genotyped them at 234 genetic markers where G and R parents had nearly-fixed (>85%) allele frequency differences, and mapped quantitative trait loci (QTLs) for all traits of interest (Fig 1B). We found a total of 7 QTLs that achieve genome-wide level of significance, including signals for cestode mass, fibrosis, granulomas, and ROS, but no QTL for cestode presence/absence or granulocyte/lymphocyte ratio (Fig 3A; Table S1). Fibrosis, for example, maps most strongly to a QTL on chromosome 2 where R alleles confer increasing fibrosis frequency (Fig. 3B). Each QTL explains a small to moderate amount of phenotypic variance in parasite resistance traits. For example, a QTL on the stickleback chromosome 12 explains 9.3% of the variation in cestode mass (Fig. 3C). Additional examples are provided in Figs. S11-12. We observe several potential cases of pleiotropy, the strongest of which involves overlapping QTLs on chromosome 15 with effects on both ROS and cestode mass (Fig. 3A).

**Fig. 3.**
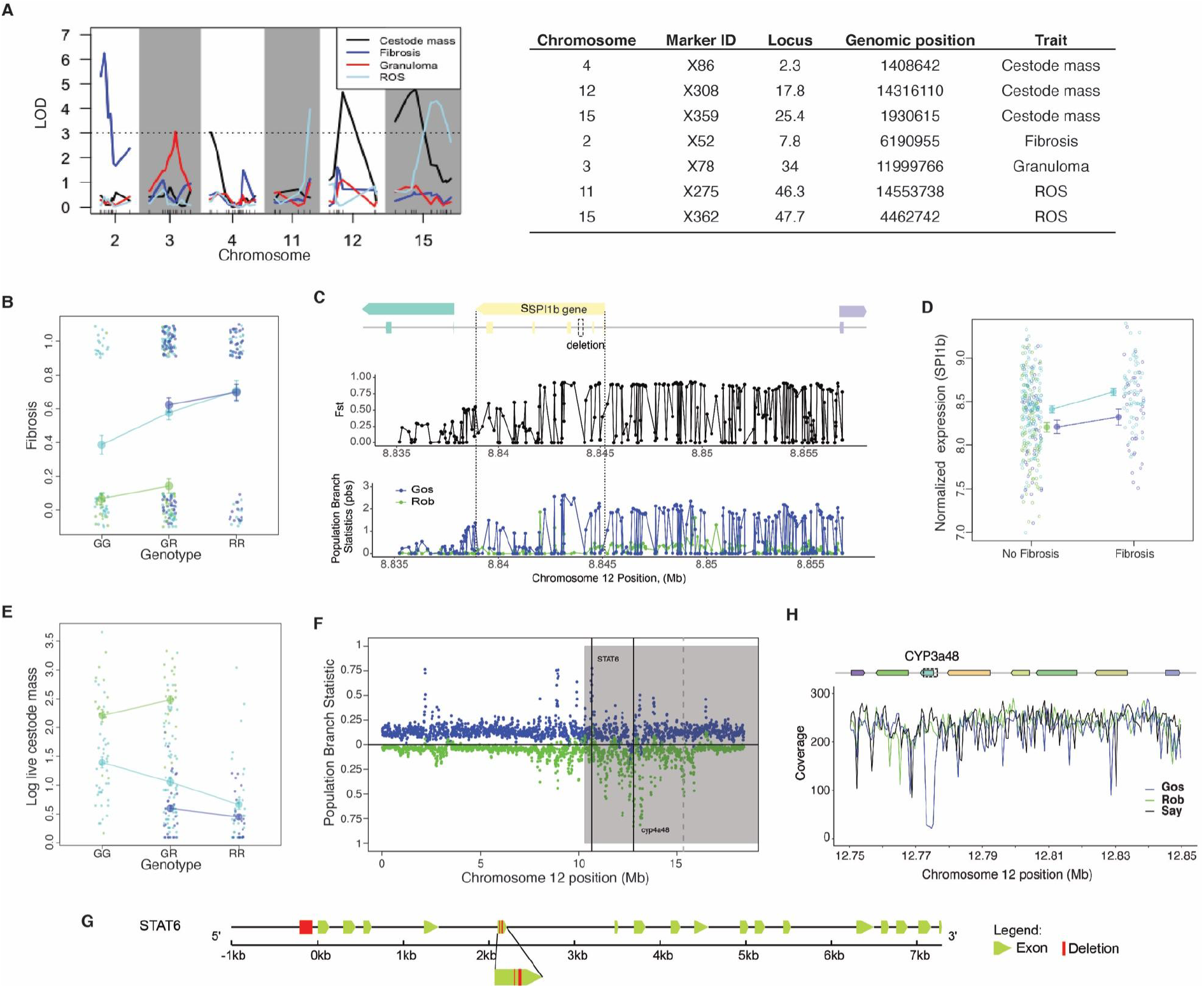
Genetic mapping and population genomic analyses of immune differences between stickleback populations. **A.** Quantitative Trait Locus (QTL) mapping identified chromosomal regions associated with R versus G differences in cestode growth, fibrosis, granuloma (without fibrosis) and ROS; here we present LOD scores for focal QTL. Tick marks on the x axis represent informative SNPs for mapping. **B.** Effect plot showing association between Chr2 marker X52 genotype and the frequency of fibrosis, for the three hybrid cross types (blue RBC, bluegreen F2 intercross, green GBC), with means and standard errors. Jitter is added to distinguish overlapping points. **C.** Effect plot of Chr12 marker X308 genotype on log cestode mass. **D.** Within the Chr2 QTL for fibrosis, PoolSeq data indicates the strongest target of divergent selection between R and G poolse is in and adjacent to the 3’ end of *PU1* gene (top subpanel is F_ST_ between R and G, bottom subpanel is the population branch statistic for G and R showing accelerated evolution is in Gosling Lake, including the fixation of a deletion within the intron containing a regulatory CTCF binding motif. **E.** The gene SPI1 (which produces PU1) is more highly expressed in fibrotic than non-fibrotic fish, controlling for infection status and cross type. **F.** Within the QTL for cestode mass and ROS on Chr12, the strongest genomic targets of selection (high population branch statistic) are tightly clustered around the genes STAT6 and cyp3a48. **G.** STAT6 contains nearly fixed deletions in Gosling Lake just 3’ to the start of the gene, and within exon 5. **H.** cyp3a48 contains a 3kb deletion within exon 2 that is fixed in G fish, as indicated by the large window of reduced coverage in G but not R or Sayward poolseq data.

To to pinpoint regions of exceptional allele frequency divergence within the broad regions of each QTL and narrow our search for candidate genes, we used whole genome sequence data from pooled population samples (PoolSeq) from Roberts and Gosling Lakes, and an outgroup (a marine anadromous population, Sayward Estuary, S), to calculate scaled population branch statistics (PBS) measures for each locus (i.e., the proportion of total allele frequency change happening in each population). We require candidate genes to fit four lines of evidence: (1) fall within a QTL for a focal trait, (2) be an exceptional target of selection within that trait (both high PBS in R or G, and high allele frequency divergence (Fst), (3) have a known effect on the focal trait, and (4) preferably be differentially expressed in transcriptomic data from pure R and G fish (Lohman et al. 2017), and in the F2 hybrids (Fuess et al. 2021). Only a few loci met three or four of the criteria. Some of the loci in question exhibit strong positive selection in the susceptible/tolerant G population, others evolved primarily in the resistant population, consistent with expectations from optimization models of quantitative immune traits (Orr 1998).

First, we focus on the QTL on chromosome 2 where R alleles confer greater fibrosis in infected fish (Fig. 3B). Within the QTL window the strongest target of divergent selection (Fst ∼ 1.0, Fig 3C, Fig.S13) is a narrow area containing the gene *SPI1b* (ENSGACT00000020522.1), which produces the transcription factor PU.1 that is known to regulate fibroblast polarization and initiation of tissue fibrosis (Wohlfahrt et al. 2019; Watt et al. 2021). This locus exhibits a 78 bp deletion in the second intron (nucleotides 8843941-8844018), which is nearly fixed in the G poolSeq data but absent in both R and ancestral S genotypes (Fig. 3C), and extensive allele frequency divergence just upstream of the coding sequence in a likely regulatory region. The Ensembl human genome browser identifies a CTCF promoter in this region, but exact homology is poor. Using 3’Tagseq data, *SPI1b* is expressed more in fibrotic compared to non-fibrotic F2 hybrid fish (Fig 3D, lfc = 0.199; padj = 0.0324), and in infected fish (lfc = 0.254, padj = 0.00364). *SPI1b* thus meets all of our criteria: it is the strongest target of selection within the QTL, expressed more in fibrotic fish, and has a known mechanistic role in polarizing fibroblasts to produce fibrosis (Wohlfahrt et al. 2019). We consider this a strong candidate gene to explain the more severe fibrotic lesions of R fish after cestode exposure. Remarkably, this gene exhibits accelerated evolution in Gosling rather than Roberts lake (Fig. 3C), implying that selection favored evolutionary loss of fibrosis in G fish. None of the other genes in this QTL meet all four criteria. Focusing on other targets of selection within the QTL (Fig.S13), the genes *TMEM39A* and *IL18R1* also have plausible links to fibrosis (Kitasato et al. 2004; Zhang et al. 2021). Some genes within the QTL are differentially expressed but not in a region of allele frequency divergence (e.g., fibronectin *fn1b*; Fig. S14).

Chromosome 12 contains a QTL that explains the largest percentage of variation in cestode mass (Fig. 3E). The two strongest targets of selection in this region entail accelerated evolution in Gosling Lake (around 10.6-10.7 Mb) and in Roberts Lake (12.78 Mb -12. 82 Mb; Fig. 3F, detailed view in Fig.S15). The former contains *STAT6*, which has been widely linked to helminth resistance in lab mice, but less clearly in rewilded lab mice (Leung et al. 2018). *STAT6* is known to play a key role in regulating inflammation and activation of alternatively activated macrophages (Czimmerer et al. 2018), and has been tied to fibrosis (Walford and Doherty 2013; Nikota et al. 2017). In G fish the *STAT6* gene has a nearly-fixed small deletion in exon 5 (Fig. 3G) that results in a frame-shift that should render mRNA isoforms with this exon non-functional, while other isoforms not located on chromosome 12 remain unchanged. The G population also carries a nearly fixed 100 bp deletion 80bp upstream of the *STAT6* start codon. The latter target of selection contains a cytochrome gene *Cyp3a48*, which exhibits a 3kb coding-sequence deletion in G but not R or S populations (Fig. 3H). *Cyp3a48* produces a protein that oxidizes nifedipine, which in turn slows development of interstitial fibrosis in renal transplant recipients (McCulloch et al. 1994), so Cyp3a48 should generally act to accelerate fibrosis; its loss in Gosling Lake fish is consistent with their slower and weaker fibrosis response, and larger cestodes.

Intriguingly, the 50kb region containing *Cyp3a48* is also among the strongest targets of selection in each of two benthic and limnetic stickleback species pairs (Haerer, Bonick and Rennison 2021), which also differ in *S*.*solidus* prevalence (MacColl 2009). The deletion seen in Gosling Lake also occurs in independently evolved Echo Lake fish, which also lack fibrosis and carry large cestodes. A third candidate gene of interest, *HNF4A*, sits on a much weaker target of selection on chromosome 12, very close to the genetic marker for our cestode size QTL (Fig. S15e). hnf4*α* (hepatic nuclear factor-4α) is a putative suppressor of fibrosis (Yeh, Bosch, and Daoud 2019).

Ingenuity Pathway Analysis of F2 hybrid transcriptomes (Fuess et al 2020) indicates that the hnf4*α* pathway is induced by infection (z-score = 0.481, *padj* = 0.032), but more strongly expressed in G fish than R fish (*z-score* = -0.8, *padj* = 5.89E-17). Higher activity of this fibrotic suppressor in G fish is consistent with their lack of fibrosis. Selection on this locus acted primarily in Roberts Lake (PBS is larger in R fish), suggesting that these fish evolved to reduce hnf4*α-*based fibrosis suppression. The other QTLs exhibit less compelling cases of divergent selection (Fig. S16).

An optimization process is expected to yield a mixture of positive and negative effect QTL (Orr 1998)within a given population. We observe such a mixture of effect directions for some traits (ROS and infection rate). Although R fish generally produce more ROS per granulocyte, QTL are not uniformly in this direction (Fig S17). On the QTL for ROS on Chr15 (P=0.0006, 11.4% of variance), R alleles confer higher ROS. The same is true for the QTL on Chr12 (P=0.002, 9.89%). However, on the Chr 11 QTL (P=0.0152, 5.8% of variance), R alleles confer lower ROS. Importantly, in all these instances the R alleles for ROS are derived (i.e., strong allele frequency difference from GOS and SAY fish), suggesting that ROB fish evolved to tightly regulate the levels of this highly reactive molecule.

## Conclusions

Intuitively, it is tempting to assume that immunity is preferable to susceptibility. When confronted with two populations that differ in resistance to a parasite, it seems reasonable that selection favored an immune gain-of-function in the more resistant population. Here, we present evidence for the contrary scenario. Some freshwater stickleback populations have evolved an effective but costly immune trait (fibrosis) that protects against severe cestode infection, but other populations lack this adaptation. Our genetic data demonstrate that this absence is not merely a failure to adapt, but a distinct strategy of infection tolerance that is favored by selection (based on population genomic divergence within trait QTL) to mitigate costs of immune response that we observe in both lab and wild fish.

The evolutionary model to explain our results is presented in Fig. 4. The lake populations of stickleback we focus on did not exist 12,000 years ago, as the lakes were covered by Pleistocene glaciers. When these glaciers melted, anadromous marine stickleback colonized freshwater to establish the permanent lake populations we study today. Using modern marine fish as a proxy for those ancestral founders (Kirch et al. 2021), we are confident that those colonists were highly susceptible to *S*.*solidus* infection (high infection success rate, fast cestode growth). Although they possess the ancient genetic pathways to initiate peritoneal fibrosis (Vrtílek and Bolnick 2021), they do not do so in response to cestode infection. Consequently, when marine stickleback are experimentally introduced into freshwater they typically exhibit exceptionally high *S*.*solidus* infection rates (e.g., ∼100% infection in Cheney Lake in Alaska, and in Heisholt Quarry on Texada Island). Given the well-understood evolutionary history of stickleback, we can therefore conclude that infection resistance in freshwater (in both Gosling and Roberts Lakes) is a recently-evolved trait. Our QTL mapping suggests that this gain of resistance is polygenic (mapping to multiple chromosomal regions), pleiotropic (several QTL affecting more than one trait). These adaptations entail a mix of positive and negative effect QTLs for some traits (e.g., ROS; Fig. S17), consistent with expectations for an optimization process rather than relentless directional selection.

**Fig. 4.**
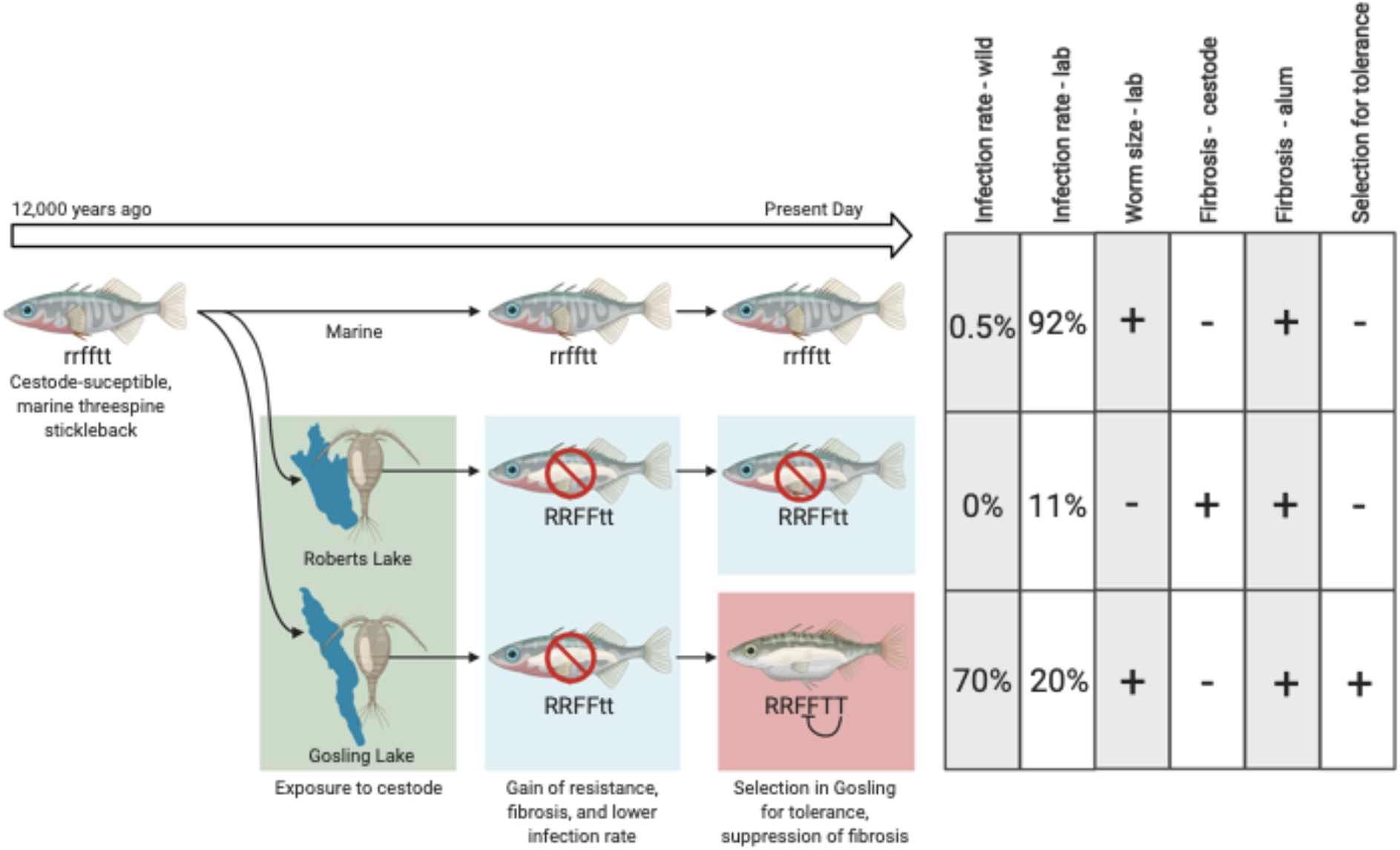
Diagram of our model of stickleback fibrosis evolution. Ancestral marine stickleback invaded freshwater, carrying alleles that confer high infection rate (r) and no fibrosis when infected (f). These genotypes persist in marine fish today, which permit high infection rates and fast cestode growth in the lab (Weber, Kalbe, et al. 2017). Newly colonized freshwater populations repeatedly evolve resistance (R, as demonstrated in Weber et al 2017) involving fibrosis (F), but the GOS population subsequently evolves tolerance (T) that suppresses fibrosis, permitting faster cestode growth. Consequently, ROB exhibits the unique phenotype (fibrosis and cestode growth suppression), but in ROB*GOS F2 hybrids the genetic difference of fibrosis maps to the tolerance loci that were subject to selection in GOS. Created with Biorender.com.

A striking element of this recently evolved immunity is our observation of extensive fibrosis in some genotypes. In humans, fibrosis is considered a severe pathology that contributes to widespread morbidity, and a large proportion of deaths in developed nations (Thannickal et al. 2014). Given this familiar pathological role, it is intriguing why some stickleback populations should have evolved such high fibrosis susceptibility and others have not. High susceptibility to fibrosis has evolved in multiple independently-derived freshwater populations of stickleback (Fig. S7), which is a classic hallmark of adaptive value. Both our experiment and field data indicate that this value comes from growth suppression, and the formation of cysts that encase and sometimes kill the cestode. In the lab, fibrosis and cestode growth were negatively correlated among cross types, and within F2 hybrids. In the field, fibrosis is associated with smaller cestodes among individuals within populations, and among populations. The consilience between our results in the lab and field observations (at multiple spatial scales) is striking evidence of a beneficial protective effect of fibrosis. The mechanism of this protection remains to be determined, but may entail reduced motility of the cestode that limits its access to nutrients or oxygen, or facilitates aggregation of immune cells on its surface.

If we simply examined the present-day phenotypic differences between populations (fibrosis and growth suppression absent in ancestor-like marine fish, and Gosling Lake), we would parsimoniously infer that the greater resistance in Roberts Lake represents the derived state. We would thus expect to see Roberts Lake fish exhibit genomic signatures of recent positive selection within the loci associated with fibrosis, and cestode growth suppression. Yet, genomic evidence is more mixed. Within the fibrosis and cestode growth QTL that differentiate R and G fish, the major targets of selection entail a mix of accelerated evolution in Roberts Lake (*TMEM39A, HNF4A*), and in Gosling Lake (*SPI1, STAT6*). *cyp3a48* is an odd case in that it sits within a window of accelerated evolution in Roberts Lake (focusing on windows of averaged PBS for SNPs), but the deletion itself is fixed exclusively in Gosling Lake indicating very localized accelerated evolution in the tolerant population. Thus, Gosling Lake fish (which are commonly infected and yet still capable of reproduction) exhibits both up-regulation of fibrosis suppression, and selective sweeps favoring deletions that are likely to disrupt function of pro-fibrotic genes. The polarization of evolutionary change clearly suggests that the differences between these two lakes entails both selection within Gosling Lake, and within Roberts Lake.

Our genetic data thus clearly support the conclusion that the phenotypic difference we study reflects divergent selection favoring tolerance (suppressing fibrosis) in Gosling Lake, and fibrosis (in Roberts Lake). The most likely model to explain our results (Fig. 4) is that during the initial colonization of freshwater, both Gosling and Roberts Lakes evolved greater resistance to S.solidus (Weber et al 2017), and that this likely entailed some gain of fibrosis in both populations. But, the substantial reproductive costs of fibrosis were untenable in Gosling Lake. We have shown here that fibrosis reduces female reproductive success in laboratory stickleback, and in wild stickleback. Indeed, the fecundity cost of fibrosis is more severe than the cestode infection (though we lack data on total fitness which includes infection-associated mortality). At present, we do not know the ecological factors that made this cost excessive in Gosling Lake (and other fibrosis-free populations), but adaptive in Roberts Lake and others like it. The difference may have to do with the relative exposure rates to S.solidus, or microbial or nutritional challenges that exacerbate the cost of fibrosis. Whatever the ecological reason, in Gosling Lake selection favored alleles that reduce the initial infection rate, as well as modifiers that mitigated these costs in the long term at the expense of allowing cestode growth. Our GxR QTL mapping cross therefore revealed QTL for the fibrosis-suppression modifiers which cause these lakes to differ phenotypically. These polymorphic loci represent an exciting opportunity to study the genetics, cell biology, and fitness consequences of fibrosis in a natural setting, to better understand the biology of this major human pathology (Wynn and Ramalingam 2012). In particular, the evolution of fibrosis suppression in Gosling Lake offers hope that with further genetic dissection we may identify genes and pathways that can be used to prevent or heal fibrosis. Transgenic and cell function analyses are needed to dissect the molecular and cellular mechanisms underlying this fibrosis polymorphism.

In 279 BC, King Pyrrhus of Epirus invaded the Italian peninsula. Despite repeatedly defeating his Roman opponents, Pyrrhus’s army suffered heavy casualties with no means of reinforcement. Despite being victorious in battle, the cost was untenable and he ended his campaign and retreated. We see a close analogy to this historical event, in the evolution of resistance by stickleback. Some populations have effectively won their co-evolutionary battle with cestodes -in wild Roberts Lake fish we see frequent fibrosis indicative of past infections that failed, but large plerceroids are entirely absent. Yet other populations have, like Pyrrhus, found the cost too great to bear and evolved a strategic retreat, with selection favoring loss of a costly immune response in favor of tolerance. This down-regulation of fibrosis is especially fascinating given the extensive pathological role of fibrosis in humans (Wynn and Ramalingam 2012), contributing to a variety of autoimmune diseases and oncogenesis. Given this familiar pathological role, the naturally-evolved (and, lost) genetic variation in fibrosis susceptibility in stickleback offers a potentially valuable model system for investigating up-and down-regulation of fibrosis by both hosts and, perhaps, by co-evolving parasites.

## Assistance

We thank Catherine Hernandez, Jessica Hernandez, Mariah Kenney, Lei Ma, Meghan Maciejewski, Kevin Pan, Jacqueline Salguero, Brandon Varela, Stijn den Haan, and Jordan Young for research assistance. The Texas Advanced Computing Center (TACC) at The University of Texas at Austin and the Center for Genome Innovation at the University of Connecticut provided HPC resources that contributed to the results reported within this paper. We thank the Genome Sequencing and Analysis Facility of the University of Texas at Austin for sequencing assistance, and the British Columbia Ministry of Environment for permitting fish collection.

## Author contributions

**Jesse Weber:** Conceptualization, methodology, investigation, analysis, visualization, writing. **Natalie Steinel:** Conceptualization, methodology, investigation, analysis, visualization, writing. **Foen Peng:** Poolseq analysis, visualization. **Kum Chuan Shim:** investigation. **Brian Lohman:** RNAseq investigation, analysis. **Lauren Fuess:** RNAseq analysis, writing. **Stephen deLisle:** field investigation of fibrosis costs, analysis. **Daniel Bolnick:** funding acquisition, conceptualization, methodology, analysis, visualization, writing, editing, supervision; **Competing interests:** Authors declare no competing interests. and **Data and materials availability:** Data and analytical code will be publicly archived on Dryad and the Short Read Archive prior to publication.

## List of Supplementary Materials

Supplemental Movie S1 Supplemental Figures S1-S17

## Methods

### Field collections

#### Variation in parasite prevalence

In 2009 we used unbaited minnow traps to collect ∼100 threespine stickleback (*Gasterosteus aculeatus*) from each of 45 lake, stream, and estuary sites on Vancouver Island. Collections were approved by the University of Texas IACUC (07-032201) and a Scientific Fish Collection Permit from the Ministry of the Environment of British Columbia (NA07-32612). Stickleback were preserved in formalin, and subsequently dissected to count *Schistocephalus solidus* cestodes and other macroparasites, to determine how infection prevalence varied among populations. Further details of the survey, including collection location coordinates and additional species of parasites see (Bolnick et al. 2020a, 2020b). For the purpose of this study we merged the resulting data with previous and subsequent years of collections from these lakes to obtain estimates of *S*.*solidus* prevalence variation among lakes spanning 2001-2015 (collections and permitting described in Bolnick 2004, Bolnick and Lau 2008, Brock et al 2017, Stuart et al 2017). We used a binomial general linear model (GLM) to test for significant between-population differences in infection prevalence, and effect of habitat type (marine, lake, stream).

To explain variation in *S*.*solidus* prevalence between lakes, we next used a binomial GLM to test for effects of log lake area, elevation, and mean log stickleback mass. For a subset of lakes (and subset of 30 individuals per lake) we also have counts of prey items from stomach contents, categorized as either benthic insect larvae or limnetic zooplankton (Bolnick et al. 2020c). In a separate binomial GLM we tested whether cestode prevalence depended on the proportion of prey that were considered benthic.

#### Covariation between fibrosis infection prevalence

In 2016, we used unbaited minnow traps to collect 431 stickleback from 13 lakes on Vancouver Island (Bob Lake, Boot Lake, Echo Lake, Farewell Lake, Frederick Lake, Gosling Lake, Muchalat Lake, Pachena Lake, Roberts Lake, and Roselle Lake). This was allowed via Scientific Fish Collection permit #NA15-217759, and UT Austin IACUC protocol AUP-2010-00024 We counted and weighed the cestodes in each individual fish, scored females them as gravid or not (based on the presence of mature eggs easily squeezed by gentle pressure), and scored fibrosis as present or absent. We tested for an association between infection (present/absent) and lake (random intercept) on the frequency of fibrosis, using a binomial GLM, including an infection*lake interaction (random slope effect). Conversely, we used a linear model to test whether log cestode mass differed between fish with versus without fibrosis, with a linear mixed model, with covariate effects of fish mass, the number of cestodes coinfecting, and a random effect of lake. Then we repeated this at the among-lake scale, using a linear model to test for a correlation between mean log cestode mass versus the frequency of fibrosis and infection intensity, with lake as the level of replication. To evaluate costs of fibrosis, we tested whether females’ reproductive state depended on the presence of a cestode, fibrosis, with lake random intercept and slope effects using a binomial GLM.

To verify these results, we revisited formalin-preserved specimens from a 2012 study of male color variation (collection permit NA12-77018A, IACUC protocol AUP-2011-00044). We dissected these fish to score the presence/absence of fibrosis and cestode infection (number and mass) from 245 stickleback from 15 additional lakes (Amor, Blackwater, Boot, Browns Bay, Cedar, Cranberry, Farewell, Gosling, Little Goose, Little Mud, Lower Campbell, Merril, Muskeg, Roberts Lakes and Sayward Estuary). Lake GPS coordinates are provided in Brock et al (2017). We again tested for an association between fibrosis and cestode infection, and between cestode mass and fibrosis, using a mixed effect GLM (*glmer* in R) with fibrosis as a function of cestode presence (fixed effect) and population serving as both a random intercept and random slope effect (e.g., a cestode*population interaction). Lastly, we replicated these results in another independent sample of 16 lakes and 16 streams (Stuart et al 2017 Nature E&E; collection permit NA19-457335, IACUC protocol 12050701). We dissected 20 formalin-preserved fish per population to score fibrosis, cestode numbers, and cestode mass, and female reproductive state (N=640 total).

We again used a mixed effect GLM to test whether fibrosis depended on cestode infection (with population as a random slope and intercept). In addition, we used a linear model to test whether log cestode mass depended on log fish mass, population, and fibrosis, focusing on the subset of populations in which both cestodes and fibrosis were prevalent (>10% of fish).

#### Male reproductive success

To assay the effects of fibrosis and male reproductive success, we snorkeled in Boot Lake (50.057 Lat, -125.533 Long) and Roselle Lake (50.522 Lag, -126.989 Long) in 2019 to hand-collect nesting male stickleback by dipnet, and non-nesting males captured by trapping outside the lek of concentrated male nests. Fish were scored for fibrosis using an ordinal 4-level system as described in Hund et al 2021, and cestodes were counted and weighed. Female gonad mass was weighed. We then used a binomial GLM to test, separately for each lake, whether nesting status of males (or gonad mass of females) differed depending on whether the fish were infected, or had fibrosis. Additional analyses and detailed methods are described in deLisle and Bolnick, (2021). These collections were approved by the British Columbia Ministry of the Environment (Scientific Fish Collecting Permit NA19-457335) and the University of Connecticut IACUC (protocol A18-008).

### Cross and experimental infection

In June 2012 we collected wild-caught stickleback from Roberts Lake (50.215 Lat, -125.542 Long) and Gosling Lake (50.068 Lat, -125.505 Long) on Vancouver Island (Scientific Fish collection permit NA12-77018 and NA12-84188). We used standard in vitro breeding methods to fertilize eggs stripped from female stickleback into petri dishes, with macerated testes from males, to make pure RxR, GxG, and reciprocal hybrid RxG and GxR families. The eggs were shipped to the University of Texas at Austin (Transfer License NA12-76852) where they were reared to maturity (The University of Texas Institutional Animal Care and Use Committee AUP-2010-00024), as described in Weber et al 2017 PNAS.

A subset of these lab-reared pure and F1 hybrid fish were experimentally exposed to *S*.*solidus* cestodes as described in detail in Weber et al. 2017. Briefly, we collected naturally infected stickleback from Gosling Lake and Echo Lake (49.987 Lat, -125.4117 Long), and shipped these to the University of Texas. Live cestodes were dissected from the euthanized fish, and size-matched pairs were placed in nylon biopsy bags in breeding media in a warm dark shaking water bath to breed. Eggs were collected from the bottom of the breeding jars after falling through the biopsy bag mesh, and stored at 4C. We hatched the eggs at 17C in 12h light per day, and fed the coracidia to *Macrocyclops albidus* copepods. We scanned individual copepods to detect infections two weeks after exposure, and fed five infected copepods to each individual fish (sample sizes and family structure described in Weber et al 2017). Exposure aquaria were checked by filtering the water after 12 hours’ feeding to confirm all copepods were consumed. Control fish included unexposed stickleback that were fed copepods without tapeworms. We euthanized stickleback 42 days post-exposure to dissect to check for infection, and to dissect out head kidneys (pronephros) for flow cytometry and transcriptomics. We did not systematically score F1 hybrid fish for fibrosis, because it was in the course of this study that we began to observe this phenotype and its association with fish genotype.

Remaining F1 stickleback were allowed to mature to reproduce. Gravid females were removed from aquaria and stripped of eggs, which we fertilized with sperm from lab-reared males to generate F2 intercross and reciprocal backcross hybrids (family cross and sample size details provided in Supplementary Table S##). These second generation hybrids were reared to maturity (∼9-15 months) and experimentally exposed to *S*.*solidus* cestodes. We exposed all available hybrid fish to copepods with tapeworms, to maximize the number of infections we could use for genetic mapping. Also, prior results showed that exposed-but-uninfected fish, and sham-exposed fish, had similar immune phenotypes (Weber et al 2017), gene expression (Lohman et al 2017), and gut microbiomes (Ling et al 2020), obviating the need for sham exposures for present purposes. Exposed fish were maintained for 42 days, then euthanized and dissected to count and weigh cestodes, and dissect out head kidneys for flow cytometry and transcriptomics (as described in Fuess et al. 2020, Lohman et al 2017). We visually scored stickleback as having fibrosis present or absent, based on whether the fish exhibited stringy sticky adhesions between the liver and intestine, or between the organs and peritoneal wall (e.g., https://www.youtube.com/watch?v=yKvcRVCSpWI). We also recorded fish sex, length, and mass. Intestines were removed and frozen for gut microbiome sequencing as described in Ling et al (2020).

### Flow cytometry -ROS production and Granulocyte:lymphocyte ratios

HK single cell suspensions were generated by manually disrupting and filtering HKs on a cell strainer (35um; Falcon 352235) in cold HK media (0.9x RPMI, 10% FBS, 100**μ**M NEAA, 100 U/ml penicillin, 100**μ**g/ml streptomycin, and 55**μ**M β-mercaptoethanol). Cell suspensions were washed once in 4ml of cold HK media (300g, 4°C, 10 min), the supernatant was removed, and the pellet resuspended in the remaining media. For each fish, ROS production was assessed by comparing DHR-123 (Sigma D1054) staining in the presence or absence of PMA (Sigma P8139) stimulation. Live HK cells were counted using a hemocytometer and 2×10^5^ HK cells were plated in 200ul of HK media, into the well of a 96 well plate. DHR-123 was added (final concentration 2**μ**g/mL) and the samples were incubated at 18C, 3%CO2 for 10 minutes. Next, PMA (final concentration 130 ng/mL) was added to the stimulated group and an equal volume of plain media was added to the unstimulated wells. Cells were incubated for an additional 20 min at 18C, 3%CO2. ROS production was determined by comparing the median fluorescence (MFI) of unstimulated and stimulated granulocytes as previously described in Weber Steinel et al., 2017. The proportion of HK granulocyte and lymphocytes was determined by flow cytometry of unstained, unstimulated HK cells via Forward and Side Scatter as previously described in Weber Steinel et al., 2017. All samples were collected on an Accuri C6 Sampler and files were analyzed using FlowJo (Treestar).

### Fibrosis scoring

Peritoneal fibrosis presence or absence was scored at the time of dissection. In healthy stickleback, the visceral organs move freely within the peritoneal cavity. Fibrosis results in the formation of adhesions between organs and/or between organs and the wall of the body cavity. Subsequently, for wild-caught fish to assay effects of fibrosis on male reproductive success we used an ordinal qualitative scoring procedure reflecting the extent and severity of fibrosis (https://www.youtube.com/watch?v=yKvcRVCSpWI).

### Analysis of infection and immune phenotypes

To evaluate the effect of overall ancestry on infection rates (cestode present/absent) in the lab infection experiments, we assigned each individual a proportion of R ancestry (e.g., 0.5 for F2 intercrosses, 0.75 in R backcrosses). We then used a binomial GLM to test whether infection depends on proportion R ancestry, sex, log mass, and room. To test effects of ancestry on cestode mass, we used a linear model with the average log cestode mass regressed on R ancestry, sex, room, log fish mass, and the number of co-infecting cestodes. Unless otherwise noted, all ANOVA tests of model significance specified type II sums of squares.

ROS burst is measured as the change in mean fluorescence intensity (MFI) of gated granulocytes with versus without PMA. We tested for genetic variation in ROS burst by regressing the change in MFI on cross, sex, log fish mass, cestode presence, and a cross*cestode interaction. The proportion of granulocytes (out of all live counted cells assigned to either granulocyte or lymphocyte gates) was also regressed on cross, sex, log fish mass, cestode presence, and cross*cestode interaction. Residuals of both regressions were checked for normality. We repeated these analyses treating genotype as an ordered quantity (proportion R ancestry) which presumes additive genetic effects. This analysis was repeated separately for infected and uninfected fish, as the latter conformed to additive genetic effects whereas the former did not.

We used a binomial GLM to test whether fibrosis (scored as present/absent) varies as a function of cross type (categorical), sex, log fish mass, cestode presence, and a cross*cestode infection. We then turned this around to ask whether cestode presence varies as a function of fibrosis, granuloma presence, cross, and cross*fibrosis or cross*granuloma interactions. To evaluate the association between granulomas and fibrosis, we used a binomial GLM to relate presence/absence of granulomas to fibrosis, cestode presence, cross type, and interactions between cross*cestode, cross*fibrosis, and fibrosis*cestode. Using linear models we tested whether cestode mass

(ln-transformed) depends on fibrosis, cross, and a fibrosis*cross interaction. We repeated the regression lumping together fibrosis and granulomas. These analyses of course were restricted to the subset of individuals with detectable cestode infections. We repeated these analyses separately for each cross type (cestode mass ∼ fibrosis) to evaluate each genotype’s trend. Next, we regressed ln cestode mass on ROS burst (delta MFI of granulocytes), with a cross and cross*ROS interaction. Finally, we used a linear model to regress ln cestode mass as a function of ROS burst, fibrosis, and cross, with all interactions, then simplified the model to drop non-significant interactions. We also tested for dependence between the explanatory variables, using a binomial GLM to test whether fibrosis depended on ROS burst (with cross and cestode presence and fish mass as covariates). As this relationship was not significant, we did not pursue this relationship further. To synthesize our evaluation of the multivariate controls on cestode mass, we conducted a path analysis (package *lavaan* in R) as illustrated in Figure 2.

We scored female stickleback as gravid if they had enlarged ovaries with full-sized clear yellowish eggs. To test the effects of infection and immune traits on female reproductive state, we used binomial GLM to test whether being gravid (T/F) depended on fibrosis, cestode presence, cross type, fish mass, and ROS burst. We repeated this analysis for wild-caught fish, testing whether females’ reproductive state depended on fibrosis, cestode presence, fish mas, and lake with lake*fibrosis and lake*infection interactions. This analysis used all 2016 wild collections as the level of replication. We also tested for a correlation between cestode prevalence and fibrosis prevalence using lake as the level of replication. We then focused within Roselle Lake, which had both moderate fibrosis and moderate cestode prevalence, to test whether female reproductive state depended on fibrosis or infection. We repeated this evaluation using a 2019 sample from Roselle Lake, using a chi square test for whether infection and fibrosis are associated.

### Masson’s Trichrome stains

Masson’s Trichrome stain was used to visualize collagen deposition within the peritoneal cavity. All visceral organs (stomach, intestine, liver, spleen) were dissected as a group from experimental fish and embedded in O.C.T (Sakura Tissue-Tek, 4583) media in cryomolds (Sakura Tissue-Tek, 4728). Tissues were flash frozen using a slurry of dry ice and 2-methylbutane, wrapped in a layer of plastic wrap and foil, and stored at -80C. Using a cryostat, 7**μ**m cross sections which traversed where the stomach, spleen, and liver are colocalized, were prepared and mounted on superfrost slides (Fisherbrand, 12-550-15). Slides were stained using a Masson Trichrome stain kit (Polysciences, 25088-1) according to the manufacturer’s instructions. Slides were imaged at 50x magnification using a Leica DM6 light microscope.

### QTL genotyping and analysis

We genotyped parental and F2-hybrid fish across several thousand variable DNA sites using a ddRADseq approach (Peterson et al 2012) that closely followed previously described protocol variants (Stuart et al 2017, Weber et al BEDASSLE 2017). Briefly, we extracted DNA from parents and hybrid fish using the Wizard SV Genomic DNA Purification kit (Promega), simultaneously digested samples with SphI and MluCI restriction enzymes (New England Biolabs), quantified concentrations with PicroGreen dsDNA assays (Life Technologies), and standardized samples to 90ng. We then simultaneously ligated samples to two types of adapters: one of 48 adapters, each containing a unique inline barcode and an SphI-compatible sticky end, as well as one universal, biotin-tagged adapter containing MluCI-compatible sticky end and an inline 8bp degenerate sequence (NNNNNIII) for PCR duplicate detection. Barcoded pools of no more than 48 samples were then run on a 2% MarkerB 100–600 bp cassette (Sage Science) to extract genomic fragments of 295–340 bp. For each sample pool we retained only biotin-tagged fragments using Dynabeads M-270 (Invitrogen), and performed six separate, twelve-cycle PCR reactions (Phusion High Fidelity PCR kit, New England Biolabs), using primers that added Flow Cell binding sequences, as well as pool-specific index-read barcodes. We sequenced our 16 total libraries across 6 lanes on a Illumina HiSeq 2500 at the Genome Sequencing and Analysis Facility, University of Texas at Austin.

To analyze sequences and generate genotypes we first demultiplexed fastq files using Stacks version 1.13 (Catchen et al. 2013), then mapped reads to the stickleback genome (BROAD version 078) using bwa-mem (version 0.7.7-r441; (Li 2013)). We used SAMtools (version 0.1.19-44428cd; (Li 2011; Li et al. 2009)) mpileup (options: -C 50, -E, -S, -D, -u, -I) to generate population-aware genotype probabilities, and BCFtools view (options: -v, -c, -g) to call individual genotypes. All sites with per individual read depths <8 were masked. We then identified sites exhibiting near-fixed SNP differences between Rob and Gos. Specifically, we sequenced 15 F1-hybrid and 8 pure strain parents, then retained only sites that were heterozygous in F1s and had at least 85% allelic divergence between pure Gos and Rob parents. Because the genome contained a number of scaffolds unlinked to the major chromosomal scaffolds, we calculated recombination frequencies and LODs between all sites in GBC and RBC fish using rQTL (V1.46-2; (Broman et al. 2003)), merged scaffolds when rf exceeded 0.7 (LOD > 3), and reordered markers using the ripple function. We identified and dropped sites that deviated from Hardy-Weinberg genotype expectations (p<0.05), and also dropped sites with excessive genotype errors based on close-range double recombination events; (Broman and Sen 2009)). We also individually scanned sites with elevated genotype error rates and masked problematic individual genotypes, the vast majority of which corresponded to homozygous calls that likely resulted from allelic dropout. We considered this a highly conservative strategy that would not lead to spurious genotype-phenotype associations but allowed us to maintain a relatively dense marker coverage. We pruned sites to retain only 1 SNP per kb (i.e., only 1 marker per RadTag), keeping the site with the highest genotype completeness across all F2 individuals. Finally, we used the fill.geno() function in rQTL to impute missing genotypes where the maximum marginal probability exceeded 0.7.

Before performing QTL scans we transformed phenotypes to approximate normality, masked outliers, and used linear models to identify which covariates to include in models, including whether fish were housed in rooms with flow-through or recirculating water systems. All significant covariates are noted in the code for building QTL models of each phenotype (see Supplemental Materials R code). We searched for QTLs using Haley-Knot regression (Haley and Knott 1992), as implemented scanone() function in rQTL, with both additive and interactive covariates when appropriate. We treated fibrosis and granuloma presence as binary traits, and analyzed both maximum mass of live cestodes per fish and ROS (i.e., granulocyte MFI) as continuous traits. In addition to scanning each cross type (i.e., GBC, RBC and intercross) separately, we also performed a cumulative scan by adding the LOD scores at all locations across the three cross types. For both individual and combined scans we performed 1,000 permutations to determine statistical significance thresholds for LOD scores. Because genome-wide significant QTLs derive their signals primarily from intercross animals, we focused only on intercross data for building multiple QTL models. We summarize QTL signals from other cross types in the supplementary code. The phenotypic effect sizes and percent variance explained by all QTL models were estimated using rQTL’s fitqtl() and refineqtl() functions.

### Population genomics analysis

We sampled 200 fish from Gosling lake, 200 fish from Robert lake, and 100 fish from Sayward Estuary in 2009. We extracted their DNA using Promega Wizard DNA extraction kit and pooled them to an equal-molar concentration for paired-end next-gen sequencing (Illumina HiSeq, at the University of Texas at Austin genome center). The three pools were sequenced to an average coverage of about 220 reads per base. We used the software BBmap (version 37.41, (Bushnell 2014)) to trim adapters (options: ktrim=r ftm=5 k=23 mink=11 hdist=2 tbo tpe), merge paired reads (default setting), and map the reads (default setting) to the BROAD version of stickleback genome (gasAcu1). Over 97% of the genome is covered in each pool. We then used the software SAMtools (version 1.9, (Li 2011; Li et al. 2009)) to filter the low quality reads (options: -q 20) and to create an mpileup file (default setting) for SNP calling. We used Popoolation2 (version 1201 (Kofler, Pandey, and Schlötterer 2011) options: --min-qual 20) for SNP calling and the R package poolfstat (Gautier et al. 2021) to calculate Fst for each SNP (options: min.rc=1, min.cov.per.pool = 10, max.cov.per.pool = 5e+03). We also calculated allele frequency differences and Population Branch Statistics for each SNP following (Yi et al. 2010). To control for variable rates of divergence among SNPs, we also calculated a scaled Population Branch Statistic (sPBS), by first calculating branch lengths between S, G, and R populations, then dividing these by the total tree length for that SNP, then calculating PBS. The sPBS for each population can then be interpreted as the proportion of the total tree length (for that SNP) that evolved in the focal population, and ranges from 0 to 1, where sPBS nearing 1 means that all of the evolutionary change occurred in the focal population (Mark Kirkpatrick, Personal Communication). Then we calculated the average PBS value within a sliding window of 20kb with a step of 10 kb through the genome. Genome-wide mean F_ST_ is 0.32 between Gosling Lake and Roberts Lake fish, 0.3 between Gosling Lake and Sayward Estuary, and 0.17 between Roberts Lake and Sayward Estuary.

### Gene expression

We collected wild adult stickleback from Roberts and Gosling Lakes in 2012 and generated pure R, G, and F1 hybrid eggs via in vitro fertilization. Fish were collected under Scientific Fish collection permit NA12-77018 and NA12-84188, shipped with Transfer License NA12-76852, and the experiment approved by The University of Texas Institutional Animal Care and Use Committee as part of AUP-2010-00024.These were raised to maturity at the University of Texas at Austin and experimentally exposed to *S*.*solidus* (5 singly infected copepods per individual) or fed uninfected copepods as a control, as described in (Weber et al. 2017). 42 days after exposure, fish were euthanized with MS-222 and head kidneys (pronephros) placed in RNAlater and frozen at -80C. RNA was subsequently extracted with an Ambion RNA extraction kit (AM1912). 3’TagSeq libraries were constructed as described in (Lohman et al. 2016, Lohman et al 2017), and sequenced at the University of Texas Genome Sequencing and Analysis Facility on a HiSeq 2500 (1×100) at a depth of ∼6.7 million raw reads per sample.

Bioinformatics steps are described in detail in (Lohman et al 2017). We used DESeq2 to test for differences in gene expression between infected versus uninfected fish (described in Lohman et al 2017), and the subjected the set of differentially expressed genes to Ingenuity Pathway Analysis to identify upstream regulatory switches that initiate cascades of differentially expressed genes.

We used the same library preparation and bioinformatic steps to obtain 3’TagSeq data on ### F2 (intercross and reciprocal backcross) fish from the QTL study described above (Fuess et al. 2021). We used DESeq2 to test for differences in gene expression between infected versus uninfected fish, and fibrotic versus non-fibrotic fish. The bulk of these analyses are reported elsewhere (Fuess et al. 2021), here we focus on reporting one specific result from Ingenuity Pathway Analysis related to an upstream regulator of fibrosis.

