## Supplemental Table and Figures for "Evolution of a costly immunity to cestode parasites is a pyrrhic victory"

**Supplementary Movie 1: https://www.youtube.com/watch?v=yKvcRVCSpWI**

**Supplementary Table 1**

| Trait | Chr | Marker | Locus(cM) | Locus(bp) | Covar | LOD | PVE |
|---|---|---|---|---|---|---|---|
| **Cestode Mass** | 4 | X86 | 2.3 | 1,408,642 | - | 3.02 | 5.8 |
| | 12 | X308 | 17.8 | 15,316,110 | - | 4.67 | 9.3 |
| | 15 | X359 | 25.4 | 1,930,615 | - | 4.76 | 9.5 |
| | 15 | X359 | 25.4 | 1,930,615 | *Room | 2.26 | 4.3 |
| | - | - | - | - | +Room | 10.32 | 22.8 |
| Full model (n = 134) | | | | | | 11.87 | 18.2 |
| **Fibrosis** | 2 | X52 | 7.8 | 6,190,955 | - | 6.60 | 13.4 |
| | 2 | X52 | 7.8 | 6,190,955 | *Granuloma | 4.49 | 8.9 |
| | - | - | - | - | +Granuloma | 6.80 | 13.9 |
| | - | - | - | - | +Room | 7.40 | 15.2 |
| Full model (n = 161) | | | | | | 15.24 | 35.3 |
| **Granuloma** | 3 | X78 | 34 | 11,999,766 | - | 3.51 | 11.3 |
| | - | - | - | - | +Fibrosis | 1.03 | 3.3 |
| Full model (n = 127, Flow-thru room only) | | | | | | 5.10 | 16.9 |
| **ROS** | 11 | X275 | 46.3 | 14,553,738 | - | 3.97 | 5.4 |
| | 15 | X362 | 47.7 | 4,462,742 | - | 4.31 | 6.4 |
| | 15 | X362 | 47.7 | 4462742 | *Room | 2.78 | 4.1 |
| | - | - | - | - | +Room | 12.37 | 19 |
| Full model (n = 237) | | | | | | 16.0 | 26.7 |

Table 1. QTL model fit summary

*=interaction covariate, +=additive covariate. All models were fitted to only F2-generation intercross fish.

**Figure S1**

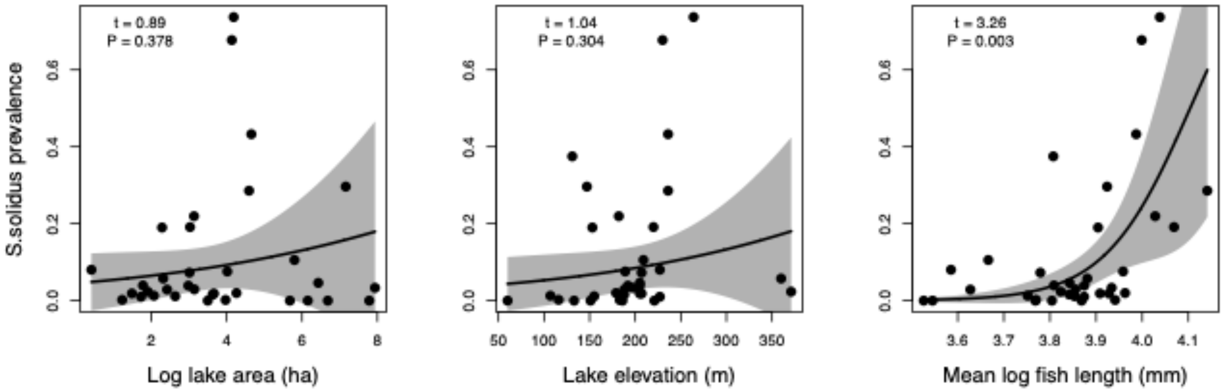


**Fig. S1.** Ecological regulation of *S.solidus* prevalence, as a function of A) log lake size, B) lake elevation, and C) mean log host mass (Binomial GLM of wild-caught lake fish, log lake area $z = 6.904$ $P < 0.00001$; elevation $z = 9.205$ $P < 0.0001$; log fish mass $z = 24.003$ $P < 0.0001$).

**Figure S2**

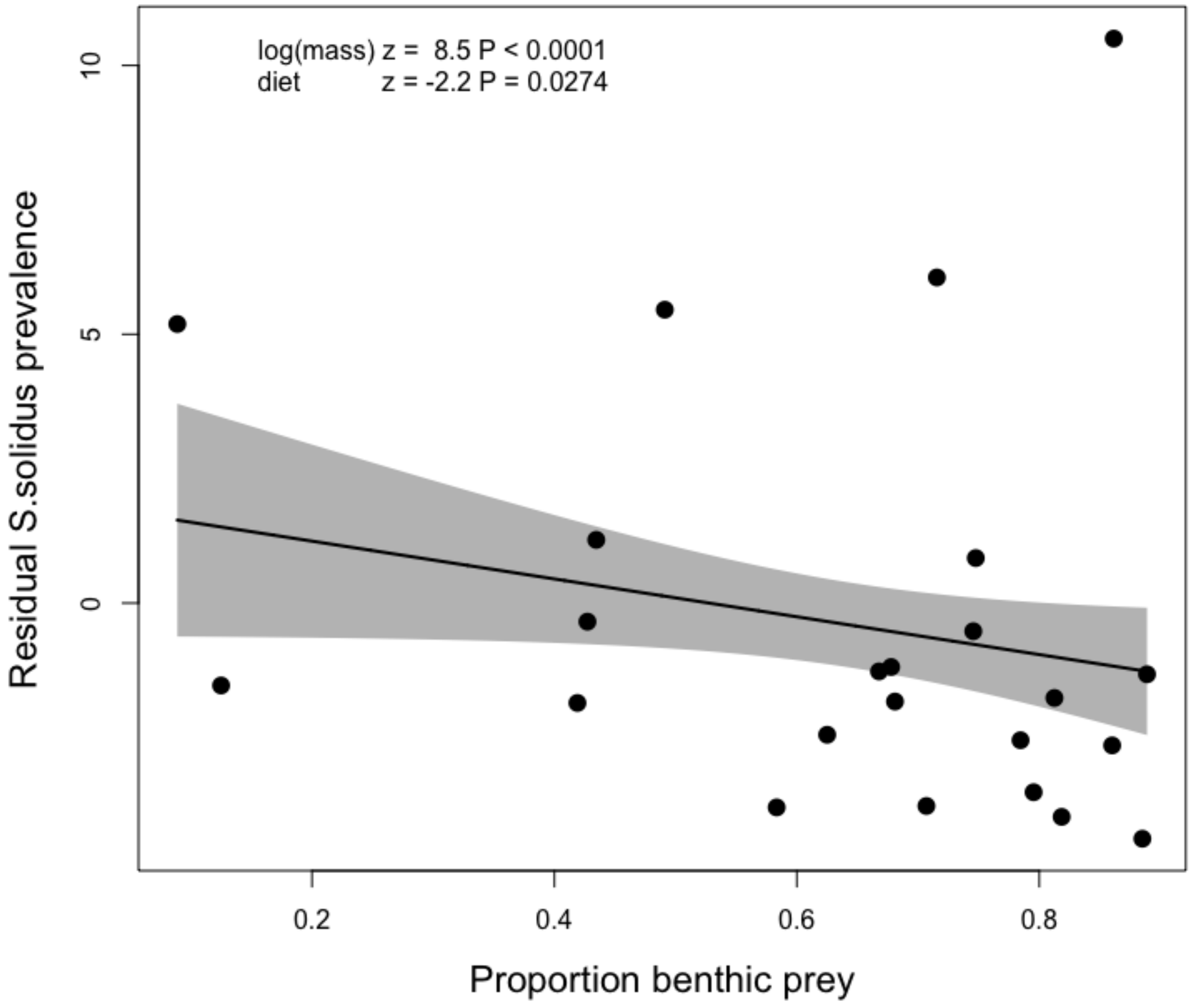


**Fig. S2.** Ecological regulation of *S.solidus* prevalence as a function of stickleback diet. We used a binomial GLM to regress cestode prevalence as a function of stickleback diet (proportion benthic prey) from wild-caught fish samples (described in Bolnick et al. 2020a,b). Controlling for lake size, elevation, and fish mass (Fig. S1), the proportion of benthic prey significantly decreases cestode prevalence (P = 0.0274), consistent with the role of limnetic prey in trophic transmission of this parasite. Here, we plot residual cestode prevalence (from a binomal GLM on log mean fish mass) as a function of the average stickleback diet estimated from stomach contents. The statistical analysis excluded Roberts and Gosling Lakes to provide independent confirmation.

**Figure S3**

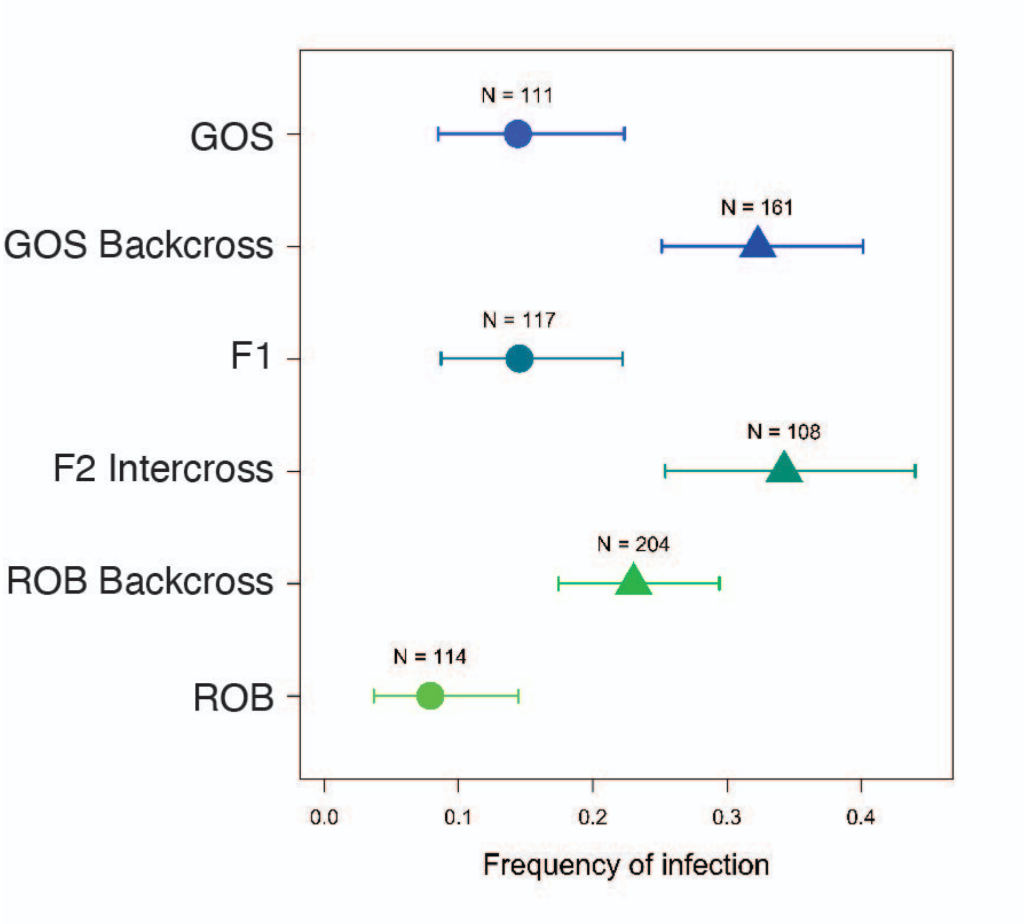


**Fig. S3.** Stickleback with greater Roberts Lake (R) ancestry have lower infection rates than those with Gosling Lake (G) ancestry. Using the F2 hybrids (F2 intercross and reciprocal backcrosses) described in this paper, Roberts ancestry significantly reduces infection rates (Z = -2.005, P = 0.045). As described in Weber et al 2017, the pure R and G fish did not differ significantly in infection rate, but the trend was in the same direction (circles denote data from that prior study). Combining these into one analysis (with study as a factor, and fish mass as a covariate), R ancestry reduces infection success ($\chi^2$ = 5.130, P = 0.0235).

**Figure S4**

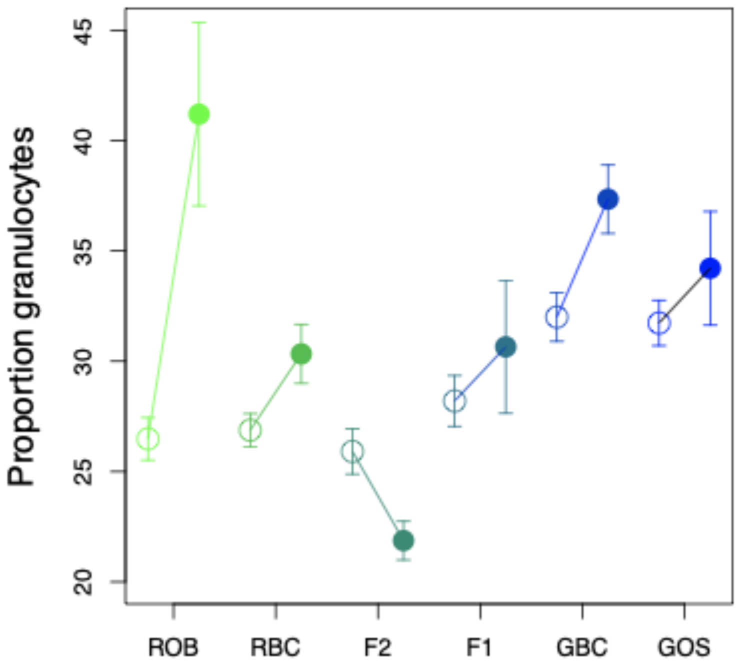


**Fig. S4.** Effect of stickleback genotype and cestode infection status on the proportion of granulocytes (versus lymphocytes) from stickleback head kidneys (pronephros), measured via flow cytometry. Filled circles are infected fish, open are uninfected. In uninfected fish, granulocytes tend to be more abundant in fish with more G ancestry (t=-4.4, P<0.0001, whereas ancestry had no linear effect in infected fish (t=-1/12, P=0.262). In all genotypes except F2 intercross hybrids, the proportion granulocytes increased in infected relative to uninfected fish (t=4.56, P<0.0001), whereas in F2 hybrids infected fish had fewer granulocytes than uninfected ones (t=-2.99, P=0.0030). As a result, there is a significant cross*infection interaction when we treat crosses as unordered factors ($F_{5,993}$=5.67, P<0.0001) in addition to significant effects of infection ($F_{1,993}$=3.90, P=0.048) and cross ($F_{5,993}$=7.24, P<0.0001).

**Figure S5**

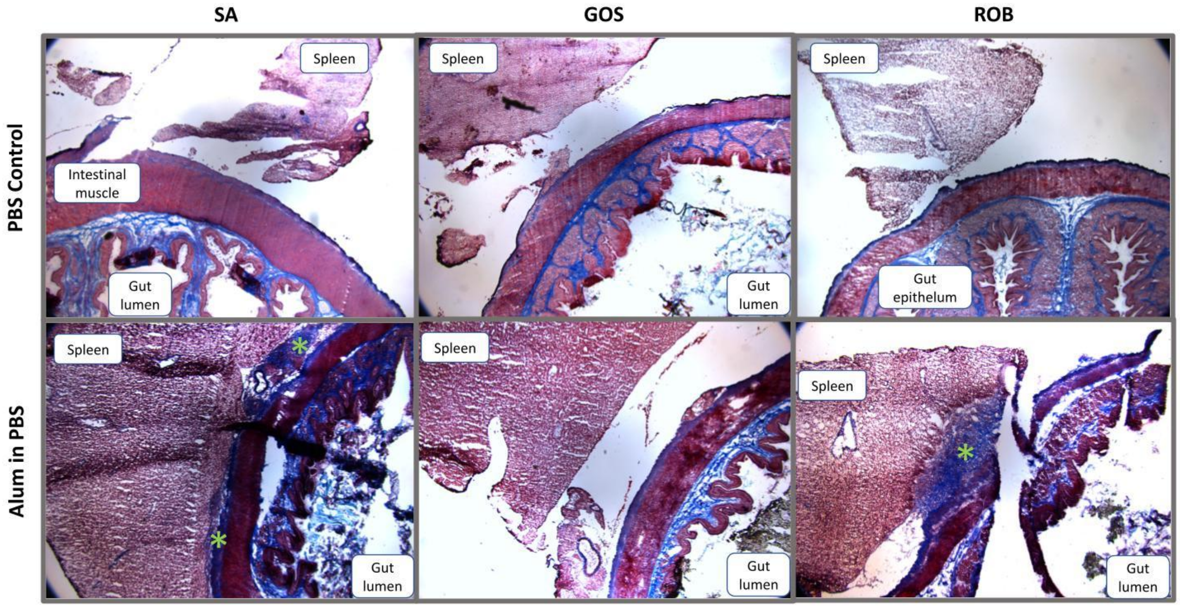


**Fig. S5.** Trichrome stain of histological sections of the gut, showing the intestinal wall, lumen, and spleen, from stickleback from the three focal populations (marine anadromous fish from Sayward, SA, and Gosling and Roberts Lakes). The fish were given intraperitoneal injections of either saline (PBS) or alum in PBS, which reproduces the peritoneal fibrosis phenotype particularly in ROB fish (bottom right). Fibrosis is stained purple and marked with green asterisks, and is absence in the PBS fish.

**Figure S6**

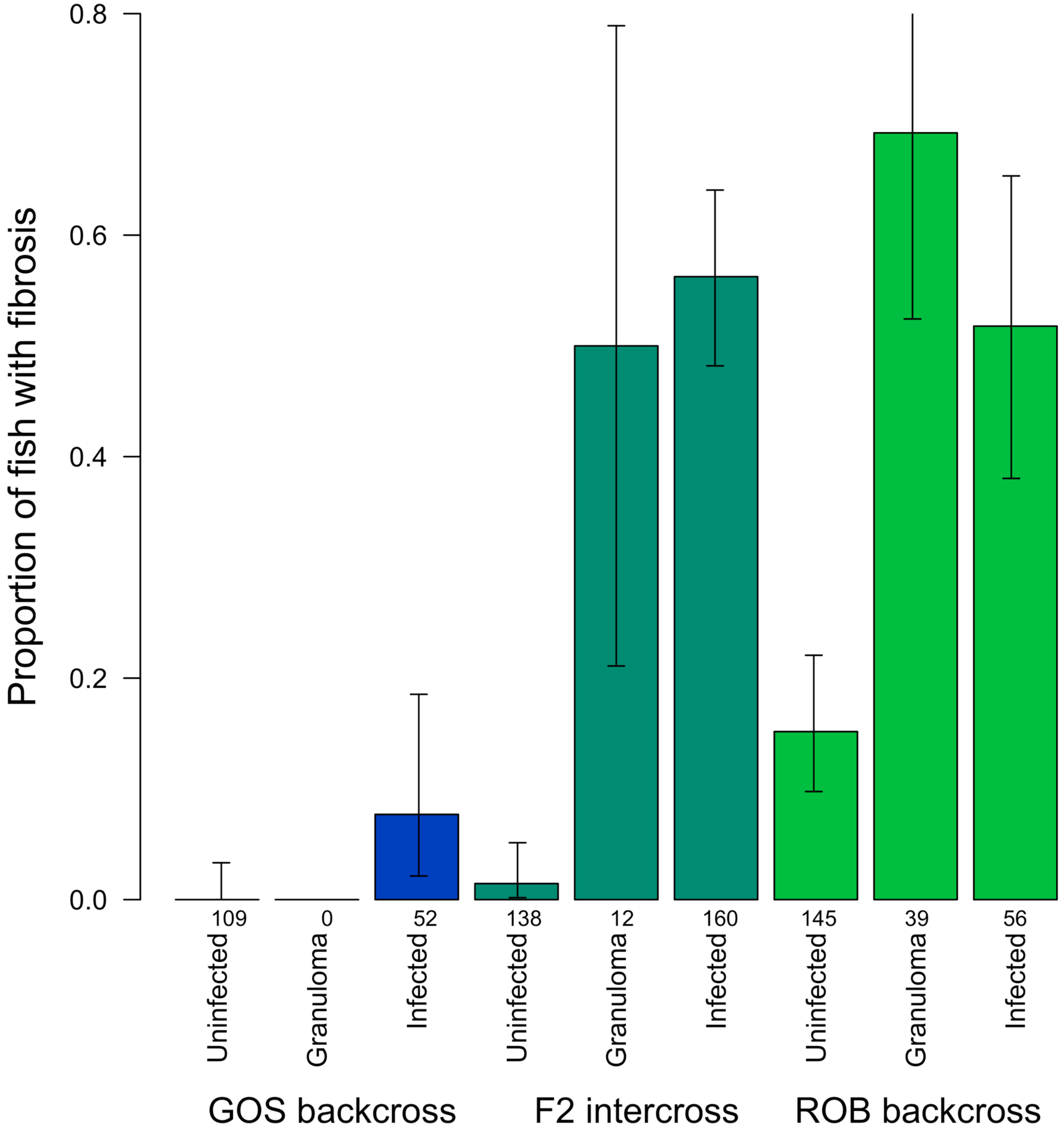


**Fig. S6.** Stickleback fibrosis frequency as a function of genotype and infection status, separating cases where the experimental infection failed (resulting in uninfected fish), or resulted in an encysted parasite (granuloma), or succeeded in establishing a freely moving peritoneal infection. ROB fish were more likely to have fibrosis even when the experimental infection had failed to colonize the body cavity. ROB fish were also relatively more likely to have an encysted rather than free-living parasite.

**Figure S7**

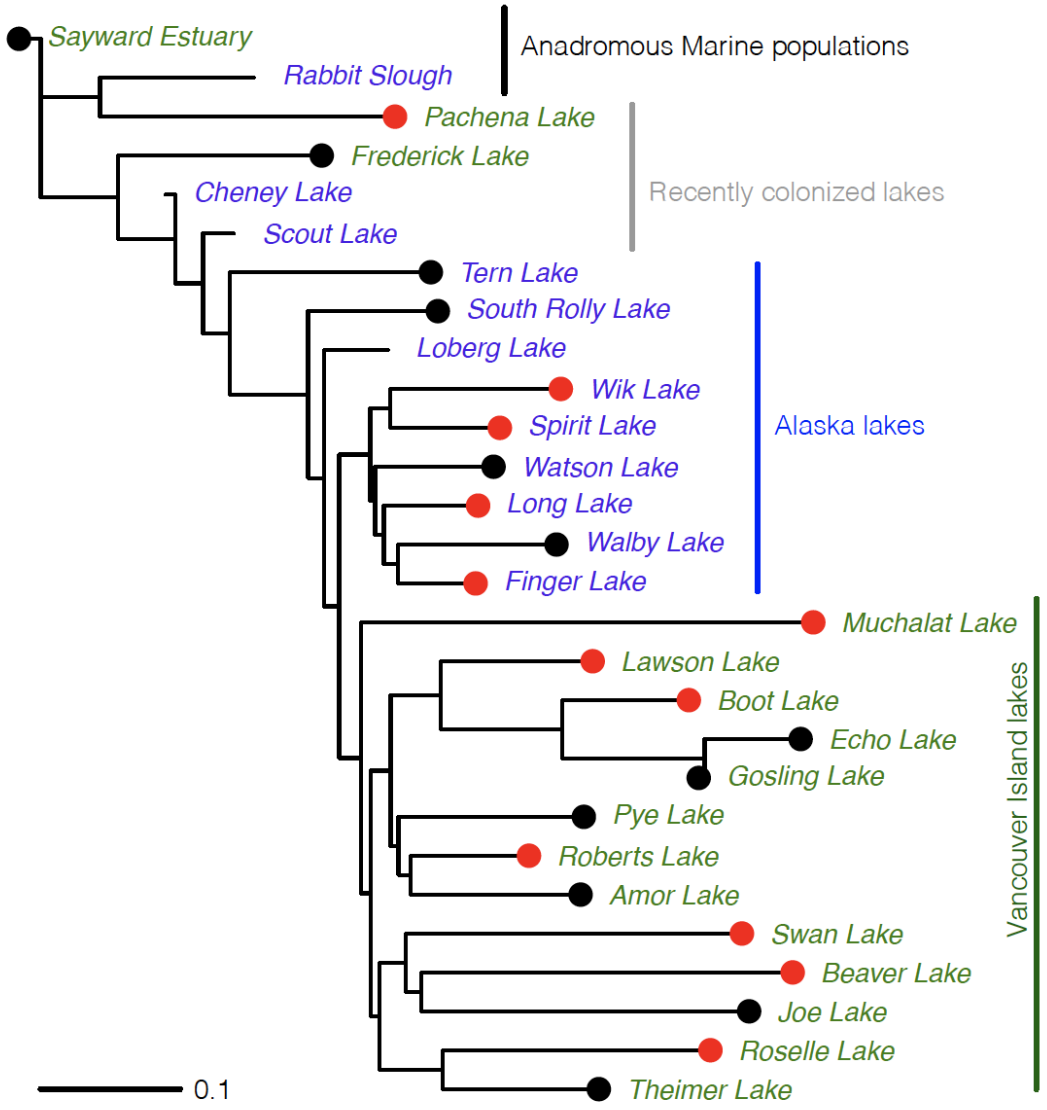


**Fig. S7.** Phylogeny of marine (Sayward Estuary and Rabbit Slough) and lake populations from Alaska (blue) and British Columbia (green), indicating populations with high versus low prevalence of fibrosis in wild-caught fish (red and black respectively). Phylogeny is based on $F_{ST}$ values (scale bar at lower left) calculated from whole-genome poolseq data of 100 fish per population, using Chromosome 21 which has few functionally important QTL and none in this study. Sayward Estuary was set as the outgroup to root the tree.

**Figure S8**

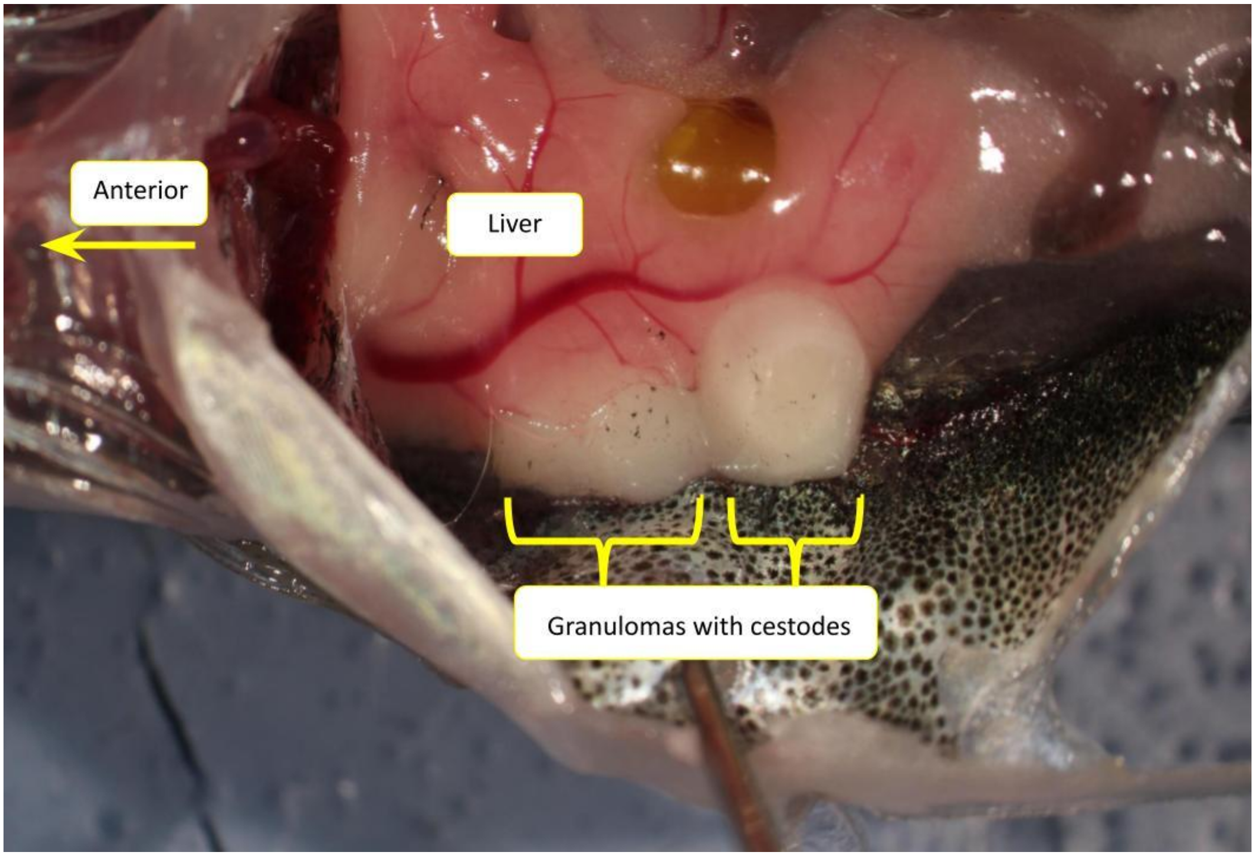


**Fig. S8.** A ventral view of a dissected stickleback (head to the left) revealing two cestodes encysted within granulomas. Both cestodes were inviable (visibly degraded and not moving), but the cysts contained more cestode RNA than host RNA.

**Figure S9**

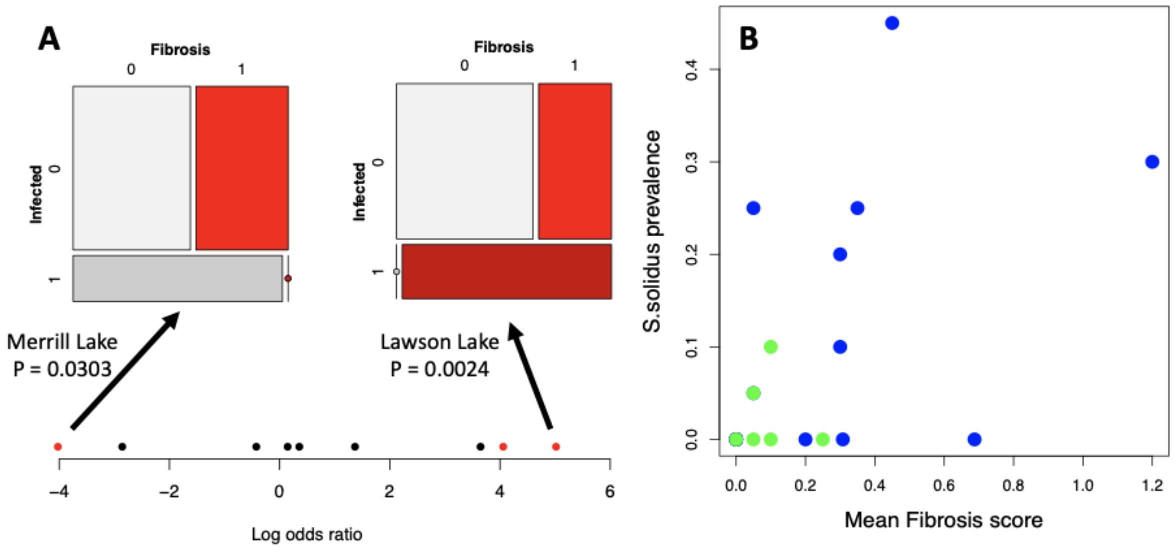


**Fig. S9.** Fibrosis covaries with cestodes in wild-caught stickleback both within and among lakes. **A)** In a survey of lakes, there were 9 populations with non-zero *S.solidus* prevalences and non-zero fibrosis, and for each lake we calculated the log odds ratio for the association between fibrosis and cestode presence in the same individuals. On average across the lakes, this association was significantly positive, but there was a significant lake*infection interaction ($\chi 2$=33.5, P<0.0001), as the infection-fibrosis association does vary among lakes. Mosaic plots from two lakes are shown. Two additional datasets from other populations and years replicated the overall positive association between infection and fibrosis at the individual fish level (Fig. S8D, $\chi 2$=4.38, P=0.0362; Fig. S8E, $\chi 2$=17.4, P<0.0001). **B)** In a survey of 16 lakes (blue) and 16 streams (green), there is a positive correlation between fibrosis severity and *S.solidus prevalence,* though as shown in Fig. 2F the cestodes tend to be smaller where fibrosis is common.

**Figure S10**

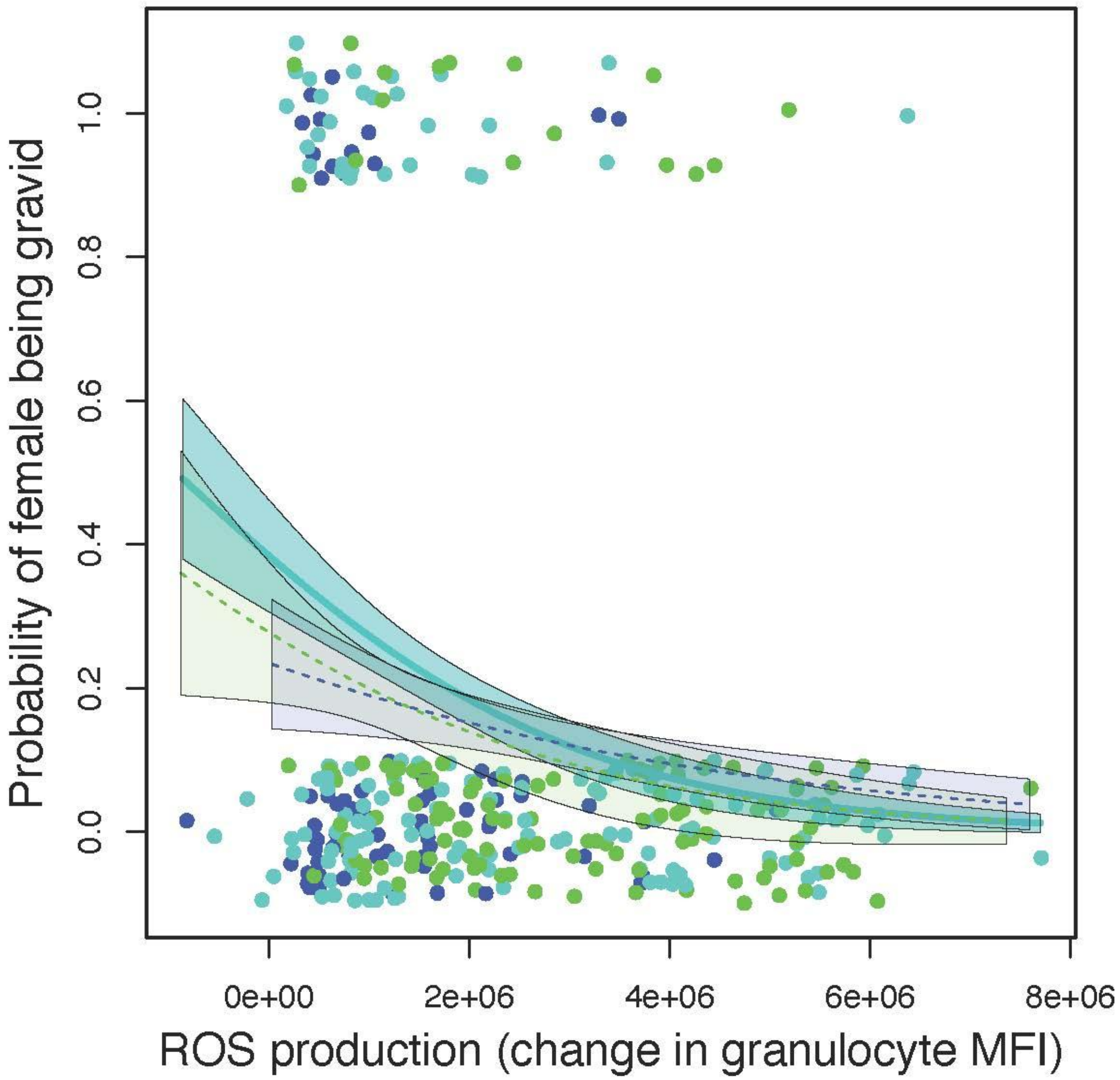


**Fig. S10.** In lab-raised F2 hybrid stickleback, female reproductive state is reduced by high ROS production by granulocytes. The effect is consistent between RBC, F2 intercross, and GBC hybrids (color coding matches other figures), though only significant within the F2 hybrids (solid trendline).

**Figure S11**

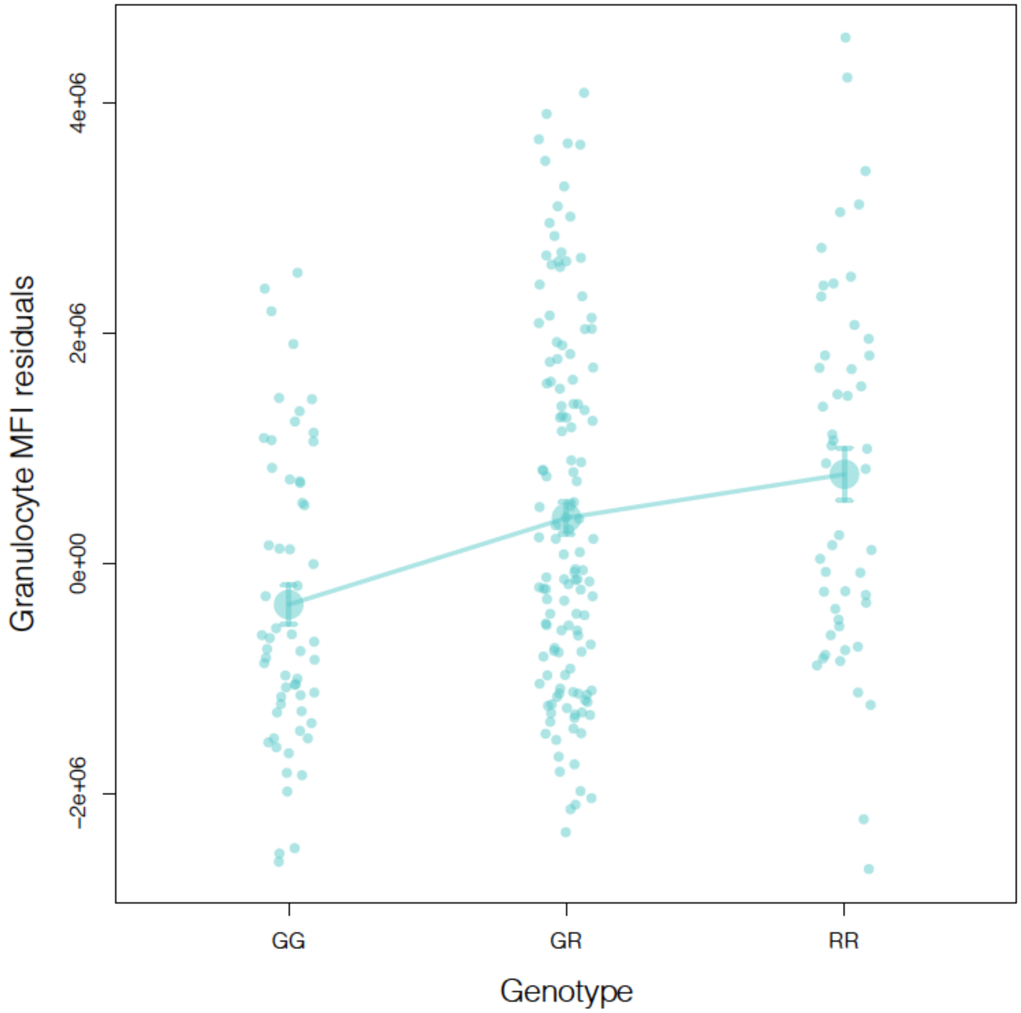


**Figure S11.** Effect plot of Chr15 marker X362 effect on ROS burst. ROS is measured by residual mean fluorescence intensity [MFI] in gated granulocytes stained with DHR and PMA stimulated prior to flow cytometry, with residuals obtained to control for daily batch effects. Here we focus on F2 intercross hybrids, showing higher ROS per granulocyte in Roberts Lake genotypes (RR) than Gosling Lake genotypes (GG). Horizontal jitter is added to distinguish points.

**Figure S12**

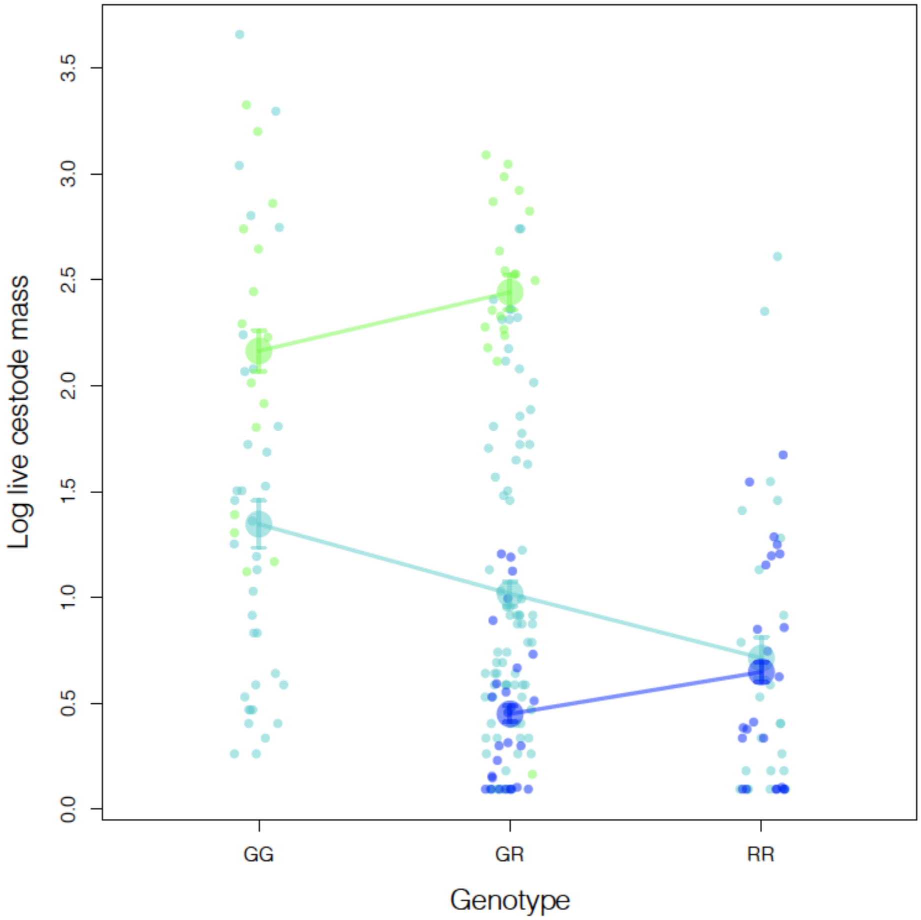


**Figure S12** Effect plot of Chr15 marker X359 effect on log cestode mass, showing three cross types (GBC in green, F2 intercross in bluegreen, and RBC in blue). Horizontal jitter is added to distinguish points.

**Figure S13**

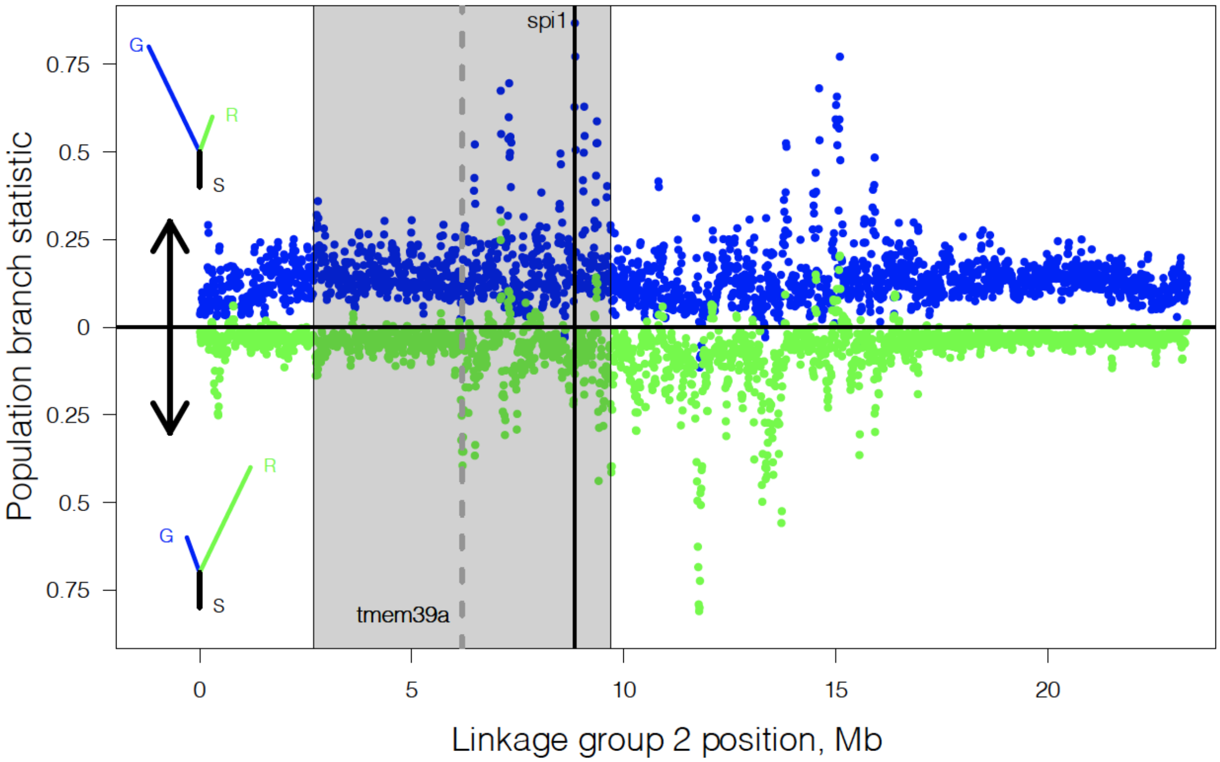


**Figure S13.** Population branch statistics for Chromosome 2, from PoolSeq data estimates of allele frequencies in R, G, and Sayward estuary as the ancestral outgroup. The plot is mirrored so the lower values (green) indicate relatively long branch lengths in Roberts Lake, and the upper values (blue) indicate relatively long branch lengths in Gosling Lake (see schematic branch lengths to the left side of the figure). The QTL window for fibrosis is shown in grey. The strongest target of selection (largest PBS) within this window contains the gene *SPI1* (aka *PU1*), as shown in greater detail in Fig. 3D.

**Figure S14.**

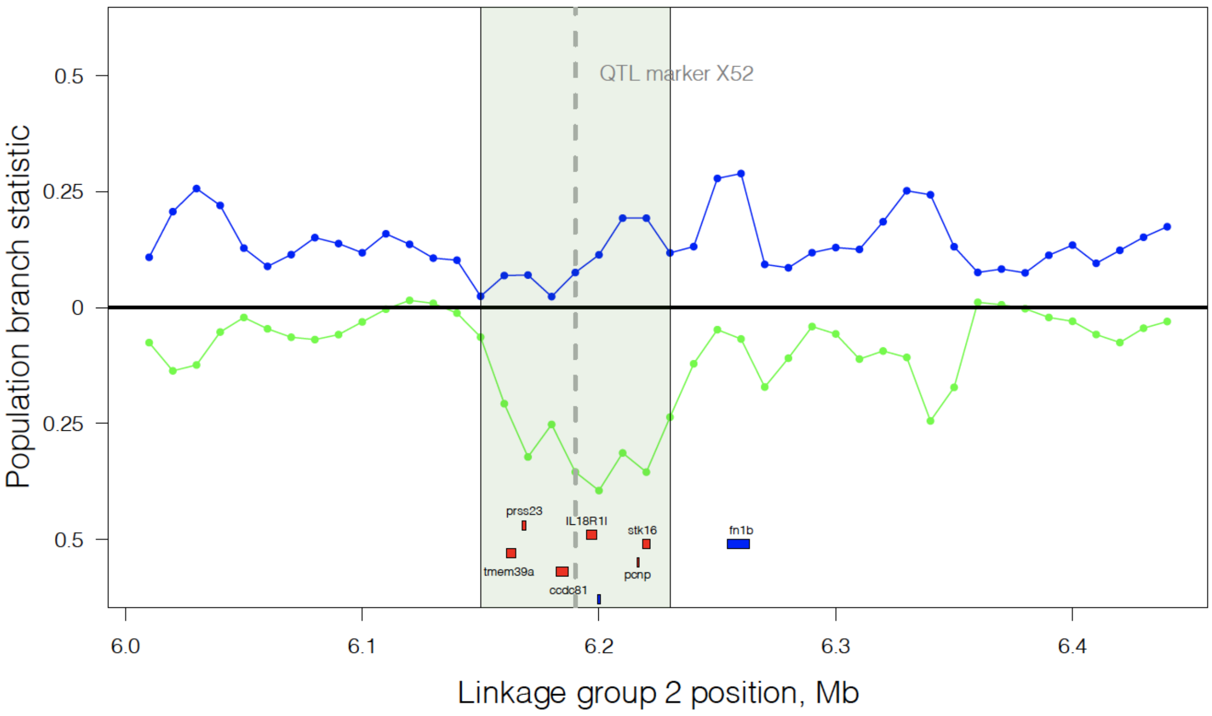
QTL marker X52
Population branch statistic
0.5
0.25
0
0.25
0.5
prss23
IL18R1l
stk16
fn1b
tmem39a
ccdc81
pcnp
6.0
6.1
6.2
6.3
6.4
Linkage group 2 position, Mb

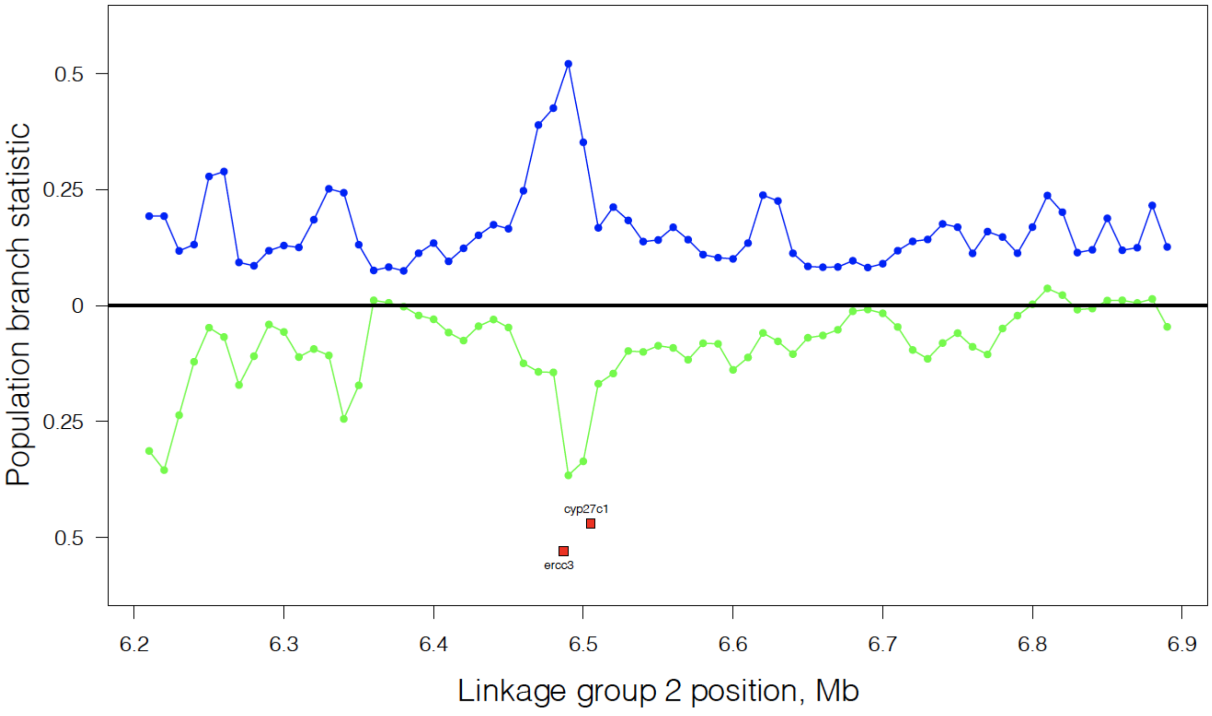
Population branch statistic
0.5
0.25
0
0.25
0.5
cyp27c1
ercc3
6.2
6.3
6.4
6.5
6.6
6.7
6.8
6.9
Linkage group 2 position, Mb

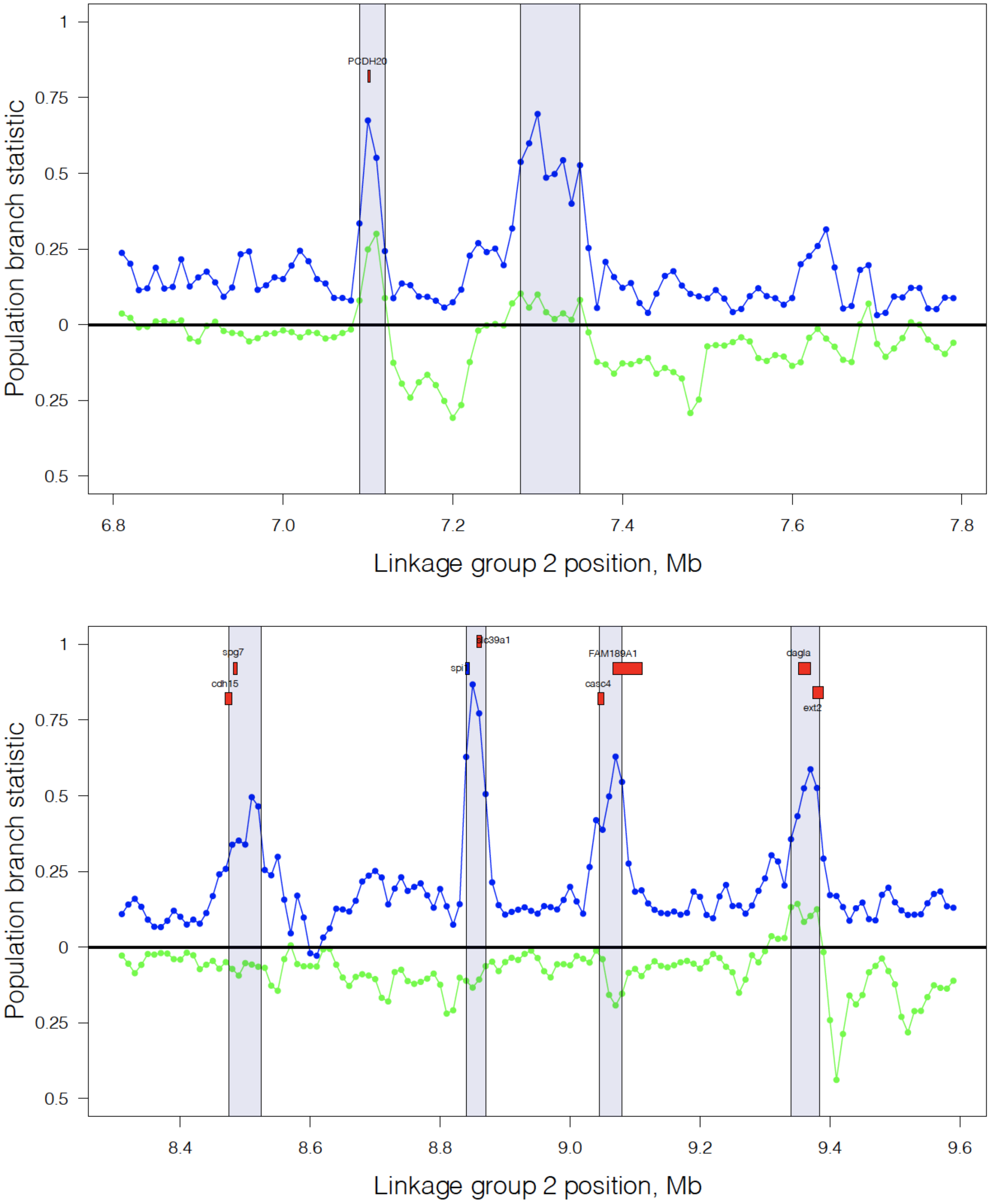


**Figure S14.** Population branch statistics for Chromosome 2, from PoolSeq data estimates of allele frequencies in R, G, and Sayward estuary as the ancestral outgroup, zooming in on peaks of accelerated evolution. Different panels focus on different peaks of selection within the QTL for fibrosis. The plot is mirrored so the lower values (green) indicate relatively long branch lengths in Roberts Lake, and the upper values (blue) indicate relatively long branch lengths in

Gosling Lake (see schematic branch lengths to the left side of Fig. S13). The QTL marker for fibrosis is shown as a dashed grey line. Genes are indicated by colored boxes with gene names. The blue gene *fn1b* (panel A) is differentially expressed as a function of genotype (LFC=0.887, Padj = 0.066), higher in R than G, but it is not in a window of allele frequency divergence. The blue gene *spi1* (panel D) is differentially expressed and a known regulator of fibrosis, and the strongest target of selection in the QTL. Blue and green shaded areas represent, respectively, accelerated evolution within Gosling and Roberts lake.

**Figure S14**

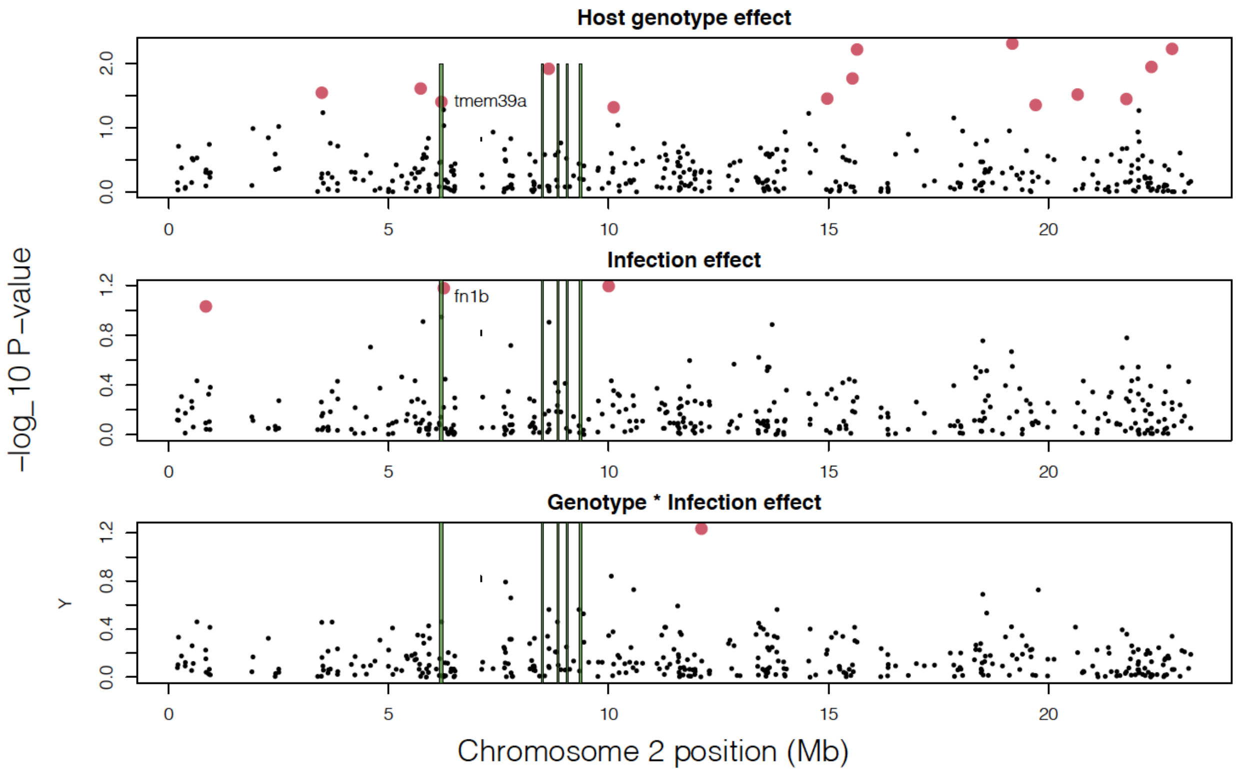


**Figure S14.** Differentially expressed genes from Lohman et al 2017, showing their position on Chromosome 2 relative to targets of selection (green bars) within the QTL for fibrosis. Separate panels convey -$\log_{10}$(adjusted P-value) for differential expression for host genotype (R versus G), infection (present versus absent) and their interaction. Red dots denote significantly differentially expressed loci within the chromosome, for the given effect. *fn1b* is differentially expressed between infected and uninfected fish, but not between genotypes, and is not within a window subject to divergent selection (e.g., no appreciable allele frequency difference between R and G). Only *spi1* is differentially expressed as a function of fibrosis (Fuess et al 2021).

**Figure S15**

**A**

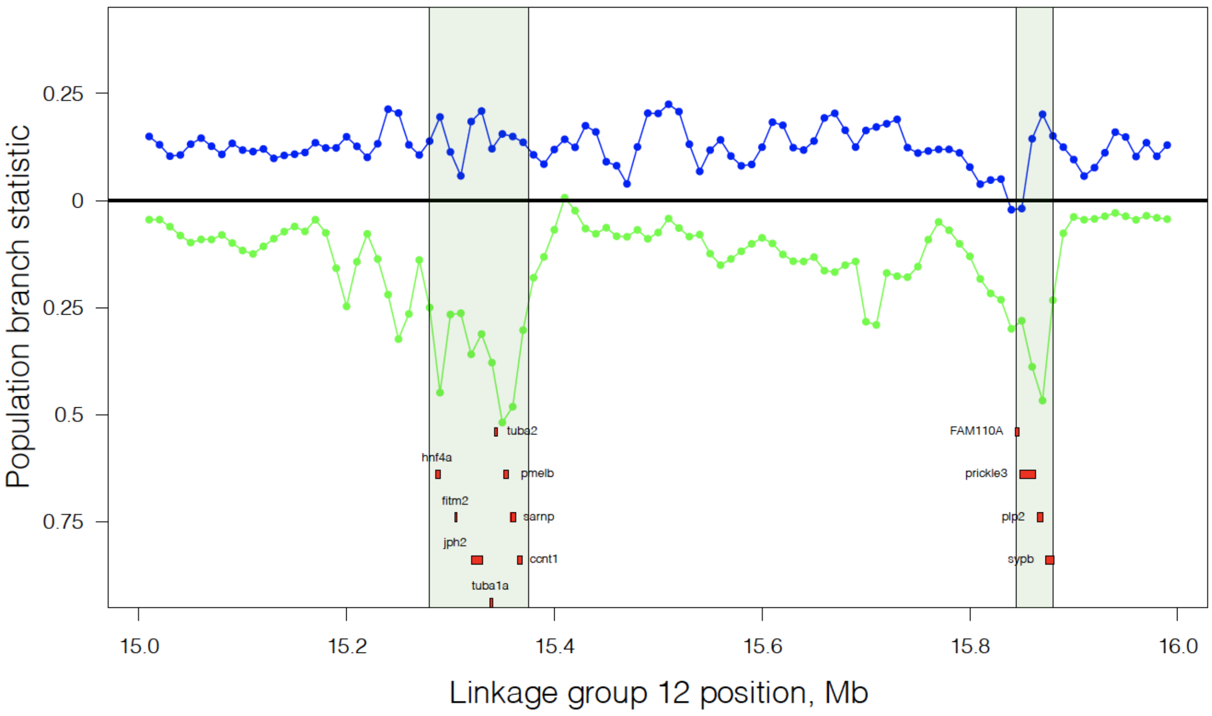


**B**

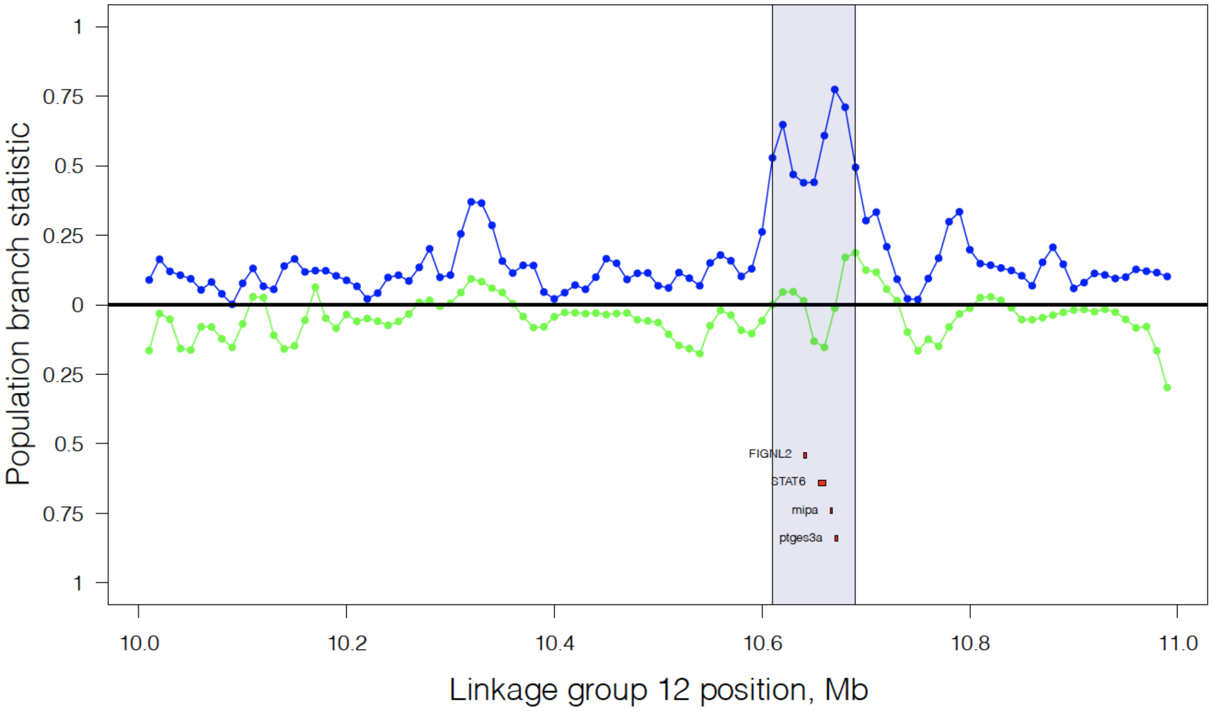


**C**

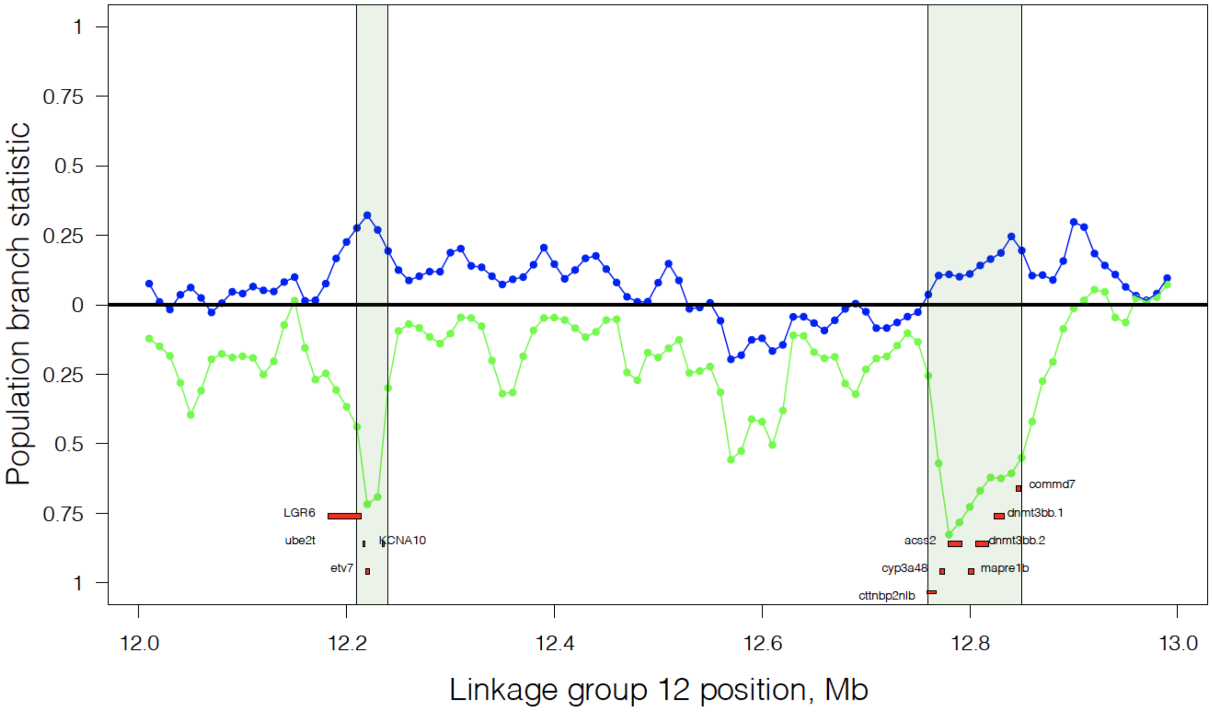
Population branch statistic
1
0.75
0.5
0.25
0
0.25
0.5
0.75
1
LGR6
ube2t
etv7
KCNA10
acss2
cyp3a48
cttnbp2nlb
commd7
dnmt3bb.1
dnmt3bb.2
mapre1b
12.0
12.2
12.4
12.6
12.8
13.0
Linkage group 12 position, Mb

**D**

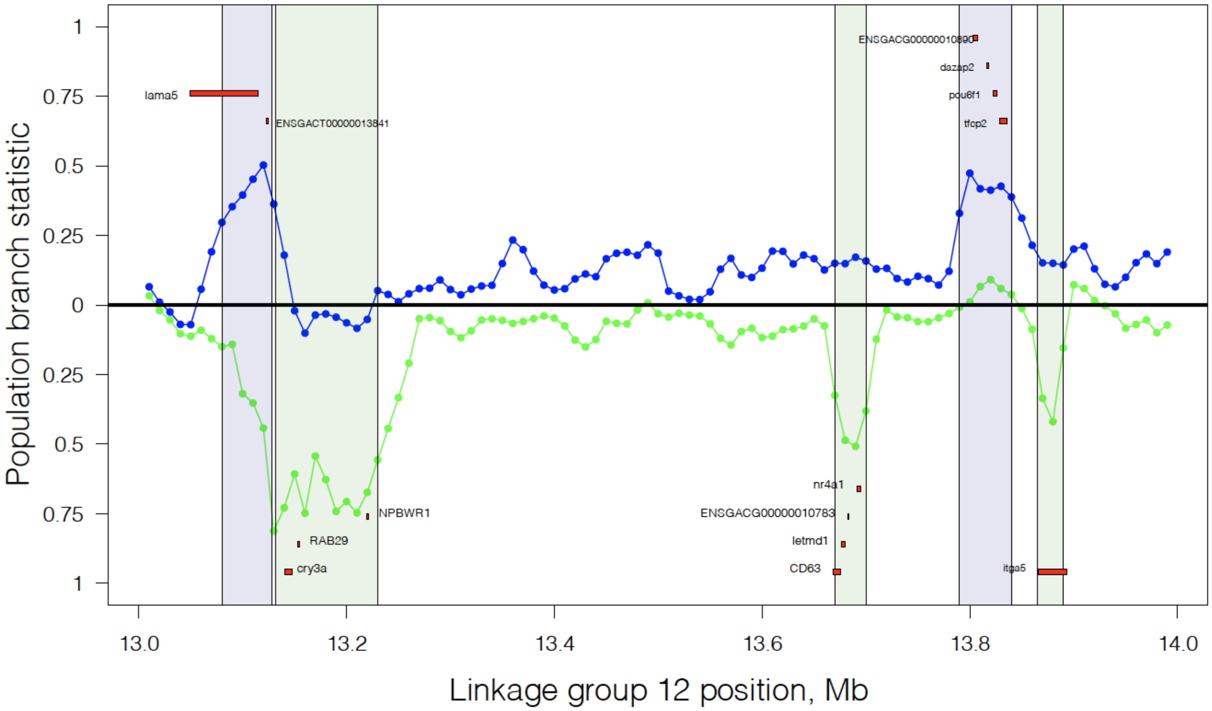
Population branch statistic
1
0.75
0.5
0.25
0
0.25
0.5
0.75
1
lama5
ENSGACT00000013841
RAB29
cry3a
NPBWR1
nr4a1
ENSGACG00000010783
letmd1
CD63
dazap2
pou6f1
tfcp2
itga5
13.0
13.2
13.4
13.6
13.8
14.0
Linkage group 12 position, Mb

**E**

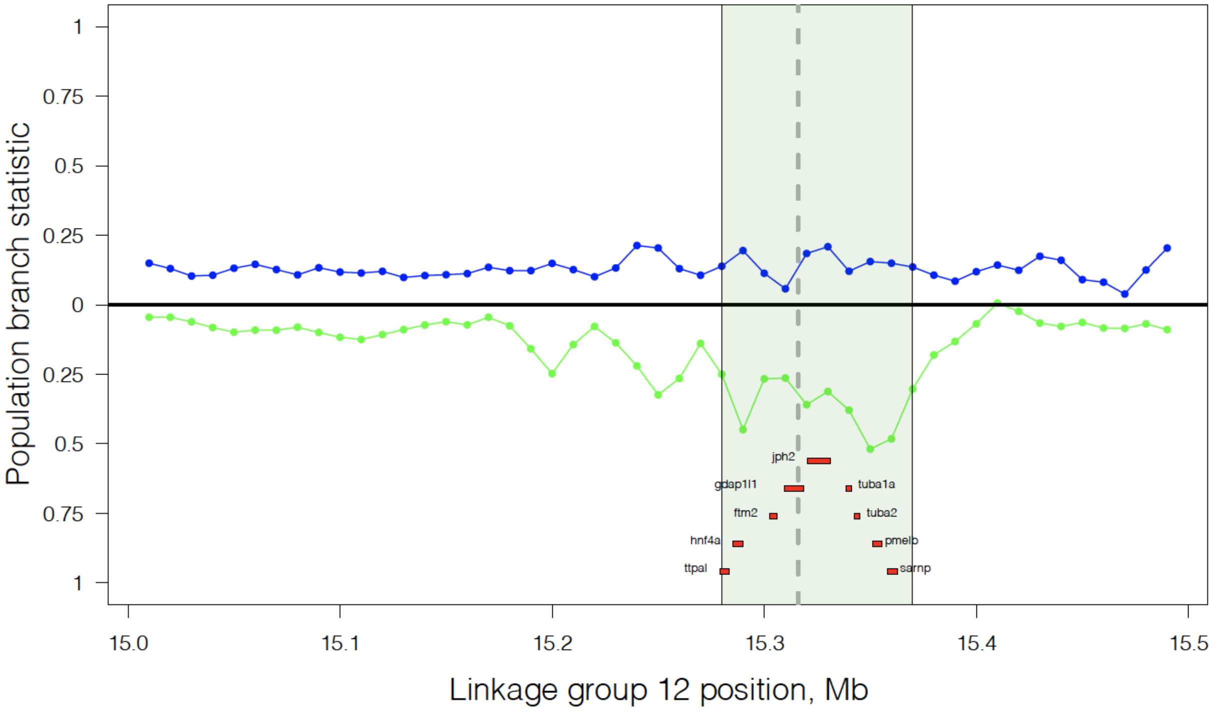


**F**

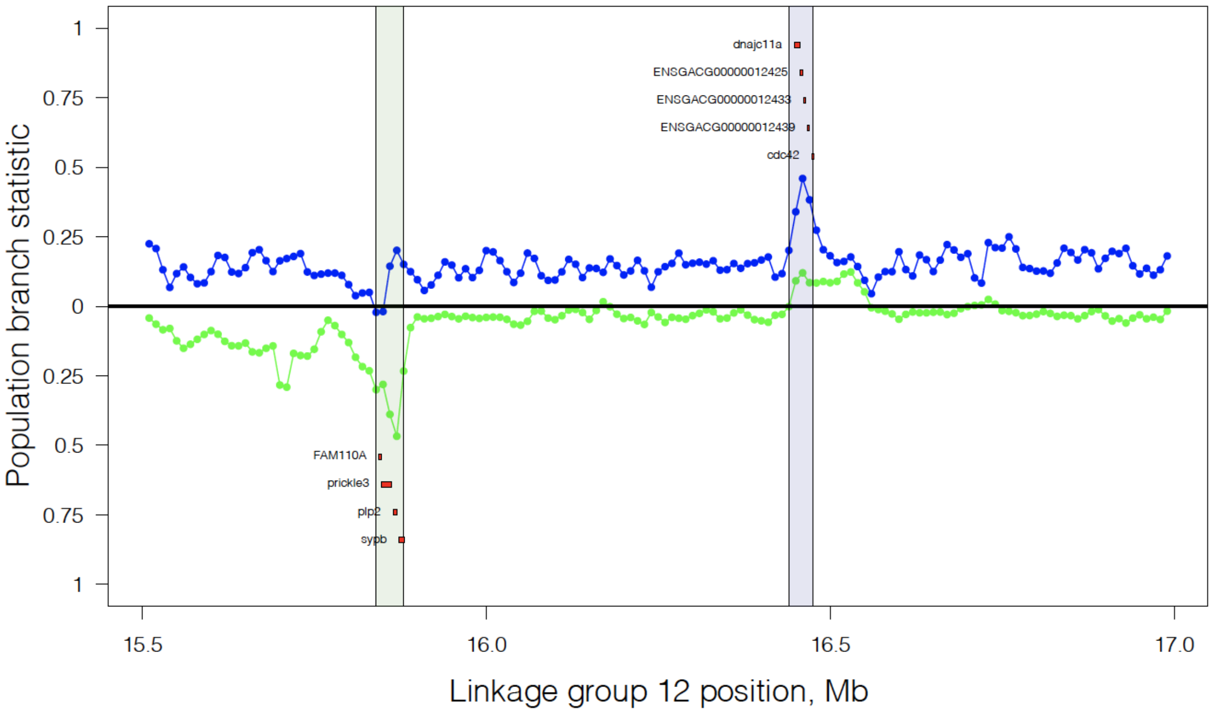


**Figure S15.** Population branch statistics for Chromosome 12 (containing the cestode mass QTL), from PoolSeq data estimates of allele frequencies in R, G, and Sayward estuary as the ancestral outgroup. The plot is mirrored so the lower values (green) indicate relatively long branch lengths in Roberts Lake, and the upper values (blue) indicate relatively long branch lengths in Gosling Lake (see schematic branch lengths in Fig. S13 for interpretation). The QTL window for fibrosis is shown in grey. Blue and green shaded areas represent, respectively, accelerated evolution within Gosling and Roberts lake. The gene *hnf4a* pathway (panel E) is the upstream regulator of a pathway that Ingenuity Pathway Analysis indicates is differentially expressed between G and R fish in response to infection.

**Figure S16.**

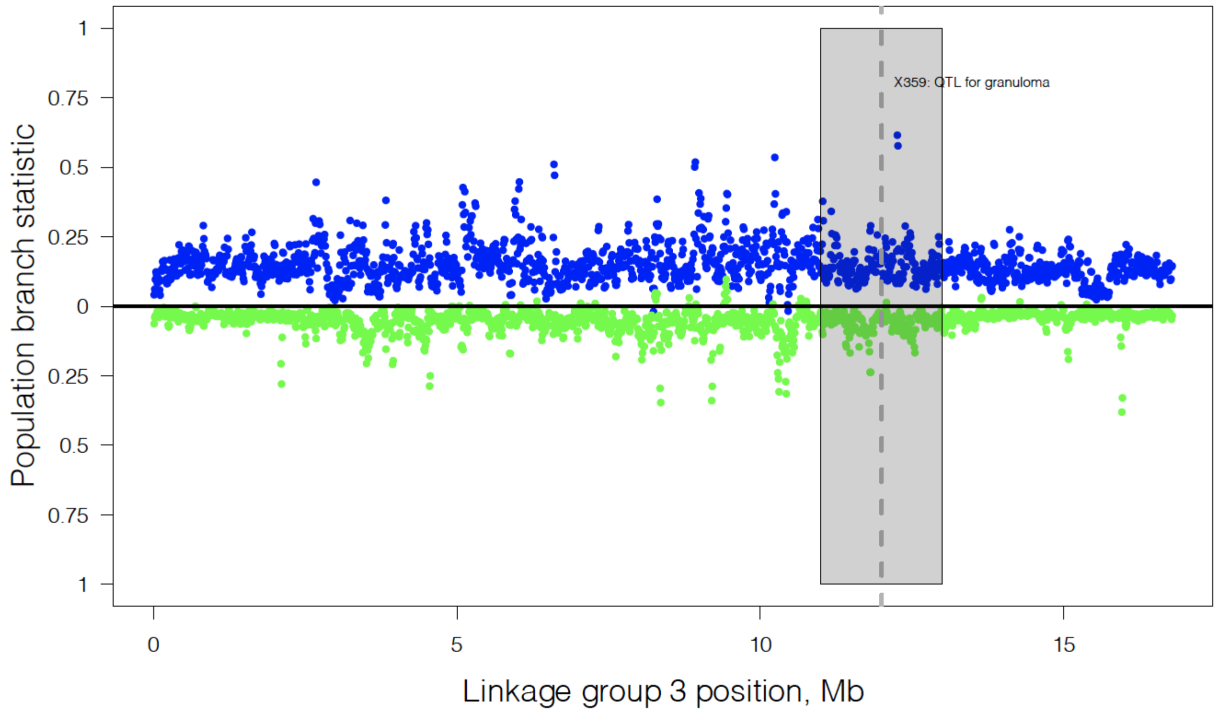
X359: QTL for granuloma
Population branch statistic
1
0.75
0.5
0.25
0
0.25
0.5
0.75
1
0
5
10
15
Linkage group 3 position, Mb

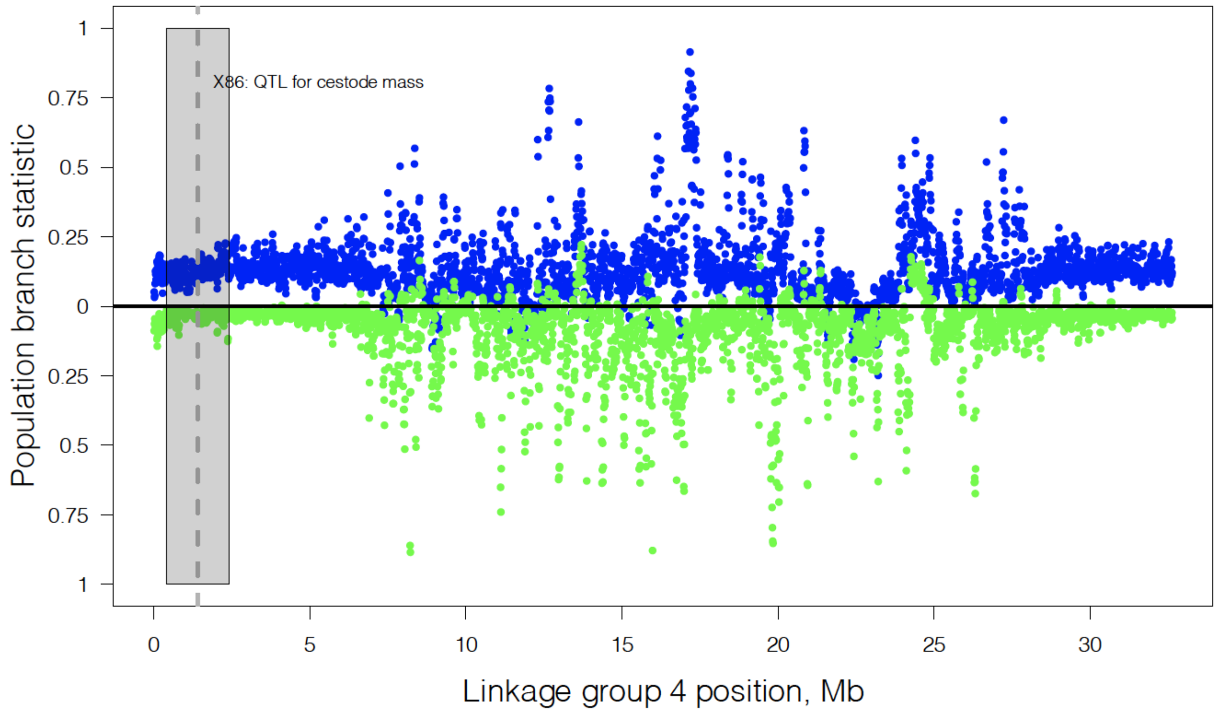
X86: QTL for cestode mass
Population branch statistic
1
0.75
0.5
0.25
0
0.25
0.5
0.75
1
0
5
10
15
20
25
30
Linkage group 4 position, Mb

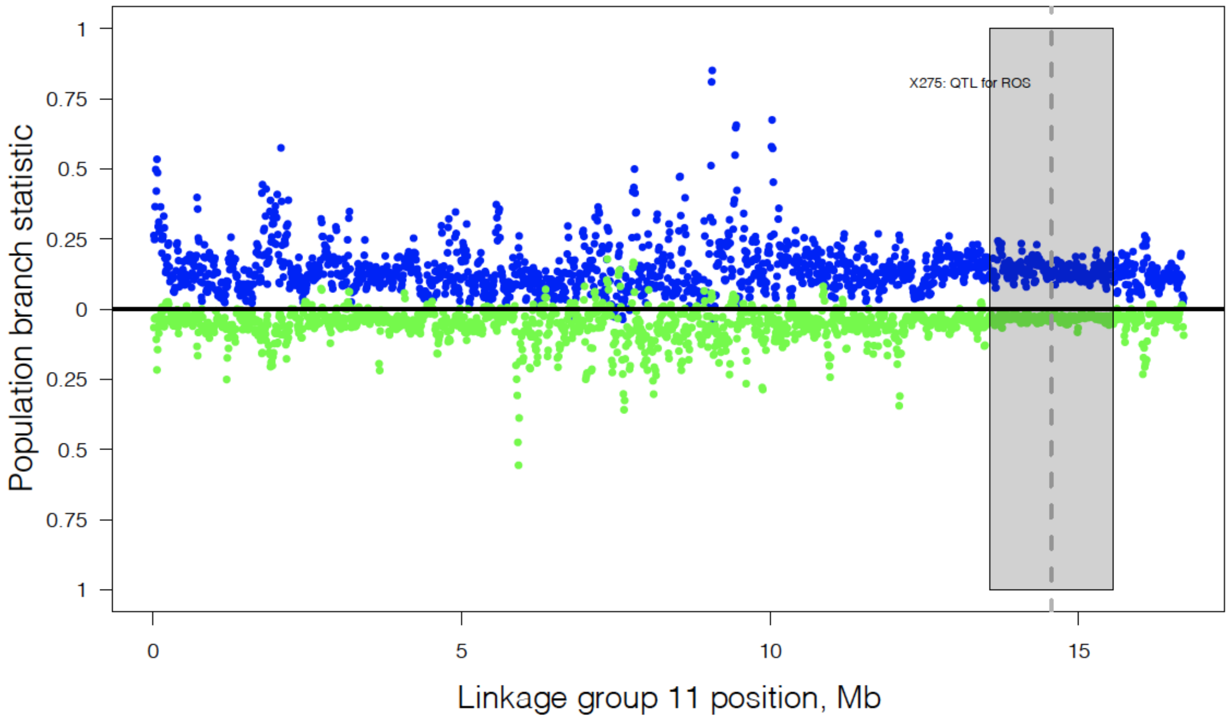


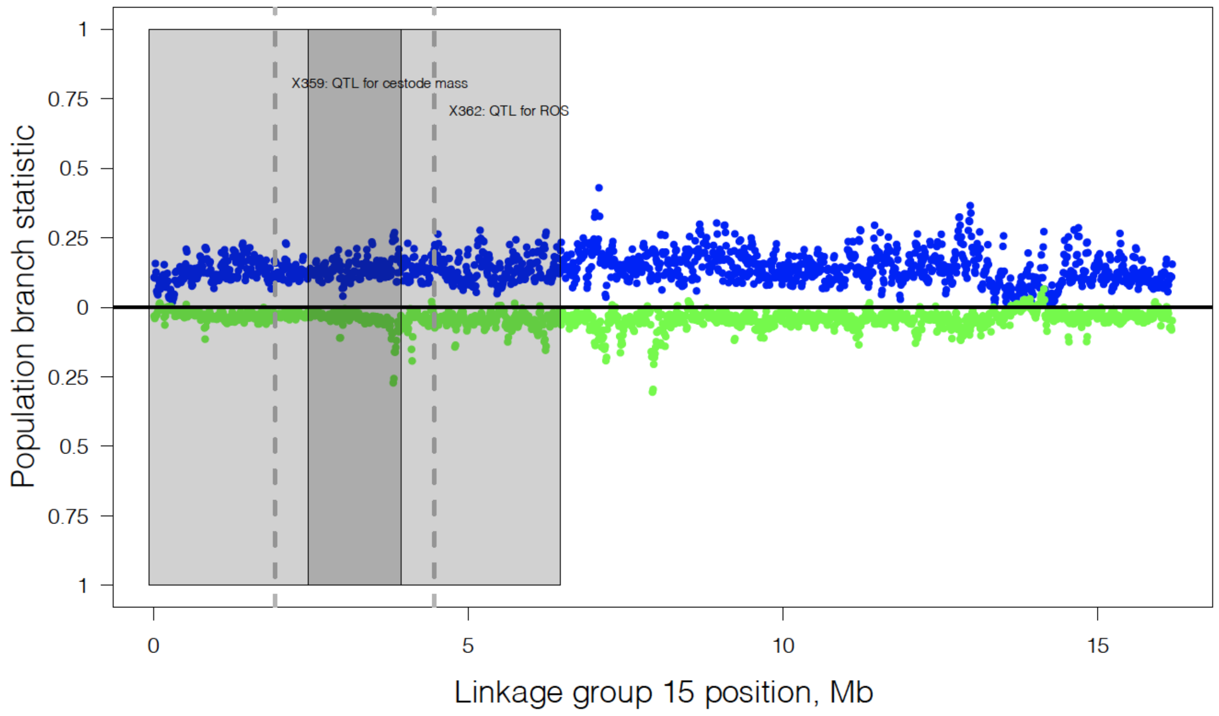


**Figure S16.** Population branch statistics for Chromosomes 3, 4, 11, and 15 (containing the cestode mass QTL), from PoolSeq data estimates of allele frequencies in R, G, and Sayward estuary as the ancestral outgroup. The plot is mirrored so the lower values (green) indicate relatively long branch lengths in Roberts Lake, and the upper values (blue) indicate relatively long branch lengths in Gosling Lake (see schematic branch lengths in Fig. S13 for interpretation). The QTL window for fibrosis is shown in grey.

**Fig S17**

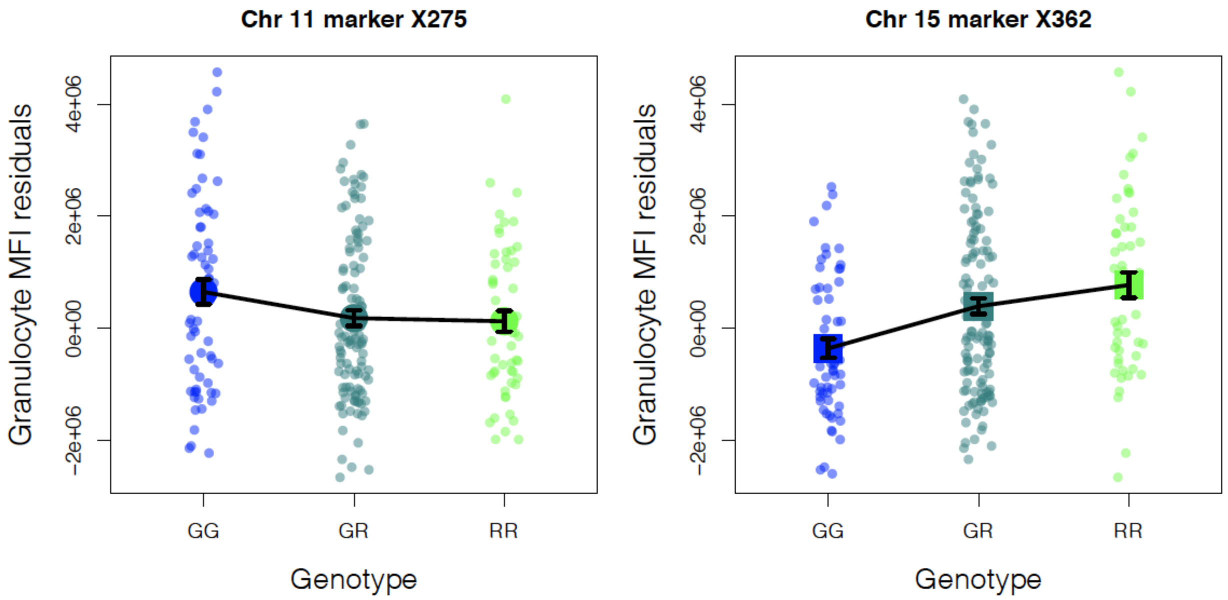


**Fig. S17.** QTL effect plots for Reactive Oxygen Species production by granulocytes, showing opposing QTL effect directions for markers on chromosomes 11 and 15, with the Roberts Lake allele decreasing ROS at the former and increasing ROS at the latter. There are three QTL supported for ROS on Chr 11, 12, and 15. Likelihood ratio tests support the retention of all three markers in an overall multilocus model, including an interaction between the Chr 11 and 12 QTL (Chr 11 F = 4.156, P = 0.0006; Chr 12 F = 3.569, P = 0.0024; Chr 15 F = 9.225, P = 0.0002; Chr 11 x Chr 12 F = 3.175 P = 0.015).

Bolnick, Daniel I., Emlyn J. Resetarits, Kimberly Ballare, Yoel E. Stuart, and William E. Stutz. 2020. "Scale-Dependent Effects of Host Patch Traits on Species Composition in a Stickleback Parasite Metacommunity." *Ecology*, September, e03181.

Boots, Michael, and Roger G. Bowers. 2004. "The Evolution of Resistance through Costly Acquired Immunity." *Proceedings. Biological Sciences / The Royal Society* 271 (1540): 715–23.

Broman, Karl W., and Saunak Sen. 2009. *A Guide to QTL Mapping with R/qtl*. Springer, New York, NY.

Broman, Karl W., Hao Wu, Saunak Sen, and Gary A. Churchill. 2003. "R/qtl: QTL Mapping in Experimental Crosses." *Bioinformatics* 19 (7): 889–90.

Bushnell, Brian. 2014. "BBMap: A Fast, Accurate, Splice-Aware Aligner." Lawrence Berkeley National Lab.(LBNL), Berkeley, CA (United States). https://www.osti.gov/servlets/purl/1241166.

Catchen, Julian, Paul A. Hohenlohe, Susan Bassham, Angel Amores, and William A. Cresko. 2013. "Stacks: An Analysis Tool Set for Population Genomics." *Molecular Ecology* 22 (11): 3124–40.

Czimmerer, Zsolt, Bence Daniel, Attila Horvath, Dominik Rückerl, Gergely Nagy, Mate Kiss, Matthew Peloquin, et al. 2018. "The Transcription Factor STAT6 Mediates Direct Repression of Inflammatory Enhancers and Limits Activation of Alternatively Polarized Macrophages." *Immunity* 48 (1): 75–90.e6.

De Lisle, Stephen P., and Daniel I. Bolnick. 2020. "Male and Female Reproductive Fitness Costs of an Immune Response in Natural Populations." https://doi.org/10.1101/2020.07.10.197210.

Ebert, Dieter, and Peter D. Fields. 2020. "Host-Parasite Co-Evolution and Its Genomic Signature." *Nature Reviews. Genetics*, August. https://doi.org/10.1038/s41576-020-0269-1.

Fuess, Lauren E., Stijn den Haan, Fei Ling, Jesse N. Weber, Natalie C. Steinel, and Daniel I. Bolnick. 2021. "Immune Gene Expression Covaries with Gut Microbiome Composition in Stickleback." *mBio* 12 (3). https://doi.org/10.1128/mBio.00145-21.

Fumagalli, Matteo, Manuela Sironi, Uberto Pozzoli, Anna Ferrer-Admetlla, Linda Pattini, and Rasmus Nielsen. 2011. "Signatures of Environmental Genetic Adaptation Pinpoint Pathogens as the Main Selective Pressure through Human Evolution." *PLoS Genetics* 7 (11): e1002355.

Gautier, Mathieu, Renaud Vitalis, Laurence Flori, and Arnaud Estoup. 2021. "F-Statistics Estimation and Admixture Graph Construction with Pool-Seq or Allele Count Data Using the R Package Poolfstat." *bioRxiv*. https://doi.org/10.1101/2021.05.28.445945.

Haley, C. S., and S. A. Knott. 1992. "A Simple Regression Method for Mapping Quantitative Trait Loci in Line Crosses Using Flanking Markers." *Heredity* 69 (4): 315–24.

Hammerschmidt, Katrin, and Joachim Kurtz. 2007. "Schistocephalus Solidus: Establishment of Tapeworms in Sticklebacks--Fast Food or Fast Lane?" *Experimental Parasitology* 116 (2):

142–49.
Hasselquist, Dennis, and Jan-Åke Nilsson. 2012. "Physiological Mechanisms Mediating Costs of Immune Responses: What Can We Learn from Studies of Birds?" *Animal Behaviour* 83 (6): 1303–12.
Heins, David C., John A. Baker, and Dillon M. Green. 2011. "Processes Influencing the Duration and Decline of Epizootics in Schistocephalus Solidus." *The Journal of Parasitology* 97 (3): 371–76.
Hund, A. K., L. E. Fuess, M. L. Kenney, and M. F. Maciejewski. 2020. "Rapid Evolution of Parasite Resistance via Improved Recognition and Accelerated Immune Activation and Deactivation." *bioRxiv*. https://www.biorxiv.org/content/10.1101/2020.07.03.186569v1.abstract.
Kirch, Melanie, Anders Romundset, M. Thomas P. Gilbert, Felicity C. Jones, and Andrew D. Foote. 2021. "Ancient and Modern Stickleback Genomes Reveal the Demographic Constraints on Adaptation." *Current Biology: CB* 31 (9): 2027–36.e8.
Kitasato, Yasuhiko, Tomoaki Hoshino, Masaki Okamoto, Seiya Kato, Yoshiro Koda, Nobuhiko Nagata, Masaharu Kinoshita, et al. 2004. "Enhanced Expression of Interleukin-18 and Its Receptor in Idiopathic Pulmonary Fibrosis." *American Journal of Respiratory Cell and Molecular Biology* 31 (6): 619–25.
Kofler, Robert, Ram Vinay Pandey, and Christian Schlötterer. 2011. "PoPoolation2: Identifying Differentiation between Populations Using Sequencing of Pooled DNA Samples (Pool-Seq)." *Bioinformatics* 27 (24): 3435–36.
Leung, Jacqueline M., Sarah A. Budischak, Hao Chung The, Christina Hansen, Rowann Bowcutt, Rebecca Neill, Mitchell Shellman, P 'ng Loke, and Andrea L. Graham. 2018. "Rapid Environmental Effects on Gut Nematode Susceptibility in Rewilded Mice." *PLoS Biology* 16 (3): e2004108.
Li, Heng. 2011. "A Statistical Framework for SNP Calling, Mutation Discovery, Association Mapping and Population Genetical Parameter Estimation from Sequencing Data." *Bioinformatics* 27 (21): 2987–93.
———. 2013. "Aligning Sequence Reads, Clone Sequences and Assembly Contigs with BWA-MEM." *arXiv [q-bio.GN]*. arXiv. http://arxiv.org/abs/1303.3997.
Li, Heng, Bob Handsaker, Alec Wysoker, Tim Fennell, Jue Ruan, Nils Homer, Gabor Marth, Goncalo Abecasis, Richard Durbin, and 1000 Genome Project Data Processing Subgroup. 2009. "The Sequence Alignment/Map Format and SAMtools." *Bioinformatics* 25 (16): 2078–79.
Little, T. J. 2002. "The Evolutionary Significance of Parasitism: Do Parasite-Driven Genetic Dynamics Occur Ex Silico?" *Journal of Evolutionary Biology* 15 (1): 1–9.
Lochmiller, Robert L., and Charlotte Deerenberg. 2000. "Trade-Offs in Evolutionary Immunology: Just What Is the Cost of Immunity?" *Oikos* 88 (1): 87–98.
Lohman, Brian K., Natalie C. Steinel, Jesse N. Weber, and Daniel I. Bolnick. 2017. "Gene Expression Contributes to the Recent Evolution of Host Resistance in a Model Host Parasite System." *Frontiers in Immunology* 8 (September): 1071.
MacColl, Andrew D. C. 2009. "Parasite Burdens Differ between Sympatric Three-Spined Stickleback Species." *Ecography* 32 (1): 153–60.
Maizels, Rick M., Adam Balic, Natalia Gomez-Escobar, Meera Nair, Matt D. Taylor, and Judith E. Allen. 2004. "Helminth Parasites--Masters of Regulation." *Immunological Reviews* 201

(1): 89–116.
McCulloch, T. A., S. J. Harper, P. K. Donnelly, J. Moorhouse, P. R. Bell, J. Walls, J. Feehally, and P. N. Furness. 1994. “Influence of Nifedipine on Interstitial Fibrosis in Renal Transplant Allografts Treated with Cyclosporin A.” *Journal of Clinical Pathology* 47 (9): 839–42.
Mittal, Manish, Mohammad Rizwan Siddiqui, Khiem Tran, Sekhar P. Reddy, and Asrar B. Malik. 2014. “Reactive Oxygen Species in Inflammation and Tissue Injury.” *Antioxidants & Redox Signaling* 20 (7): 1126–67.
Most, Peter J. van der, Berber de Jong, Henk K. Parmentier, and Simon Verhulst. 2011. “Trade‐off between Growth and Immune Function: A Meta‐analysis of Selection Experiments: Trade-off between Growth and Immune Function.” *Functional Ecology* 25 (1): 74–80.
Nikota, Jake, Allyson Banville, Laura Rose Goodwin, Dongmei Wu, Andrew Williams, Carole Lynn Yauk, Håkan Wallin, Ulla Vogel, and Sabina Halappanavar. 2017. “Stat-6 Signaling Pathway and Not Interleukin-1 Mediates Multi-Walled Carbon Nanotube-Induced Lung Fibrosis in Mice: Insights from an Adverse Outcome Pathway Framework.” *Particle and Fibre Toxicology* 14 (1): 37.
Nuismer, Scott L. 2017. “Rethinking Conventional Wisdom: Are Locally Adapted Parasites Ahead in the Coevolutionary Race?” *The American Naturalist* 190 (4): 584–93.
Nystrand, M., and D. K. Dowling. 2020. “Effects of Immune Challenge on Expression of Life-History and Immune Trait Expression in Sexually Reproducing Metazoans-a Meta-Analysis.” *BMC Biology* 18 (1): 135.
Orr, H. Allen. 1998. “THE POPULATION GENETICS OF ADAPTATION: THE DISTRIBUTION OF FACTORS FIXED DURING ADAPTIVE EVOLUTION.” *Evolution; International Journal of Organic Evolution* 52 (4): 935–49.
Rauw, Wendy M. 2012. “Immune Response from a Resource Allocation Perspective.” *Frontiers in Genetics* 3 (December): 267.
Schmid-Hempel, Paul. 2008. “Parasite Immune Evasion: A Momentous Molecular War.” *Trends in Ecology & Evolution* 23 (6): 318–26.
Thannickal, Victor J., Yong Zhou, Amit Gaggar, and Steven R. Duncan. 2014. “Fibrosis: Ultimate and Proximate Causes.” *The Journal of Clinical Investigation* 124 (11): 4673–77.
Tierney, J. F., and D. W. Crompton. 1992. “Infectivity of Plerocercoids of Schistocephalus Solidus (Cestoda: Ligulidae) and Fecundity of the Adults in an Experimental Definitive Host, Gallus Gallus.” *The Journal of Parasitology* 78 (6): 1049–54.
Tschirren, Barbara, and Heinz Richner. 2006. “Parasites Shape the Optimal Investment in Immunity.” *Proceedings. Biological Sciences / The Royal Society* 273 (1595): 1773–77.
T. Schultz, Eric, Michelle Topper, and David C. Heins. 2006. “Decreased Reproductive Investment of Female Threespine Stickleback Gasterosteus Aculeatus Infected with the Cestode Schistocephalus Solidus: Parasite Adaptation, Host Adaptation, or Side Effect?” *Oikos* 114 (2): 303–10.
Urban, Mark C., Reinhard Bürger, and Daniel I. Bolnick. 2013. “Asymmetric Selection and the Evolution of Extraordinary Defences.” *Nature Communications* 4 (1): 2085.
Viney, Mark E., Eleanor M. Riley, and Katherine L. Buchanan. 2005. “Optimal Immune Responses: Immunocompetence Revisited.” *Trends in Ecology & Evolution* 20 (12): 665–69.
Vrtílek, Milan, and Daniel I. Bolnick. 2021. “Phylogenetically Conserved Peritoneal Fibrosis

Response to an Immunologic Adjuvant in Ray-Finned Fishes." *bioRxiv*. https://doi.org/10.1101/2020.07.08.191601.
Walford, Hannah H., and Taylor A. Doherty. 2013. "STAT6 and Lung Inflammation." *JAK-STAT* 2 (4): e25301.
Watt, Stephen, Louella Vasquez, Klaudia Walter, Alice L. Mann, Kousik Kundu, Lu Chen, Ying Sims, et al. 2021. "Genetic Perturbation of PU.1 Binding and Chromatin Looping at Neutrophil Enhancers Associates with Autoimmune Disease." *Nature Communications* 12 (1): 2298.
Weber, Jesse N., Martin Kalbe, Kum Chuan Shim, Noémie I. Erin, Natalie C. Steinel, Lei Ma, and Daniel I. Bolnick. 2017. "Resist Globally, Infect Locally: A Transcontinental Test of Adaptation by Stickleback and Their Tapeworm Parasite." *The American Naturalist* 189 (1): 43–57.
Weber, Jesse N., Natalie C. Steinel, Kum Chuan Shim, and Daniel I. Bolnick. 2017. "Recent Evolution of Extreme Cestode Growth Suppression by a Vertebrate Host." *Proceedings of the National Academy of Sciences of the United States of America* 114 (25): 6575–80.
Wohlfahrt, Thomas, Simon Rauber, Steffen Uebe, Markus Luber, Alina Soare, Arif Ekici, Stefanie Weber, et al. 2019. "PU.1 Controls Fibroblast Polarization and Tissue Fibrosis." *Nature* 566 (7744): 344–49.
Wynn, Thomas A., and Thirumalai R. Ramalingam. 2012. "Mechanisms of Fibrosis: Therapeutic Translation for Fibrotic Disease." *Nature Medicine* 18 (7): 1028–40.
Yeh, Matthew M., Dustin E. Bosch, and Sayed S. Daoud. 2019. "Role of Hepatocyte Nuclear Factor 4-Alpha in Gastrointestinal and Liver Diseases." *World Journal of Gastroenterology: WJG* 25 (30): 4074–91.
Yi, Xin, Yu Liang, Emilia Huerta-Sanchez, Xin Jin, Zha Xi Ping Cuo, John E. Pool, Xun Xu, et al. 2010. "Sequencing of 50 Human Exomes Reveals Adaptation to High Altitude." *Science* 329 (5987): 75–78.
Zhang, Zhe, Shuo Luo, Guilherme Oliveira Barbosa, Meirong Bai, Thomas B. Kornberg, and Dengke K. Ma. 2021. "The Conserved Transmembrane Protein TMEM-39 Coordinates with COPII to Promote Collagen Secretion and Regulate ER Stress Response." *PLoS Genetics* 17 (2): e1009317.
